# Extra-hematopoietic immunomodulatory role of the SCID-susceptibility gene DOCK-2 identified by stepwise maturation of human iPSCs into clonogenic mesodermal stromal progenitors

**DOI:** 10.1101/2020.07.07.192385

**Authors:** Cornelia Scharler, Rodolphe Poupardin, Patricia Peking, Martin Wolf, Gabriele Brachtl, Laurence Daheron, Katharina Schallmoser, Karsten Jürchott, Harald Stachelscheid, Hans-Dieter Volk, Dirk Strunk

**Affiliations:** Cell Therapy Institute, Spinal Cord Injury and Tissue Regeneration Center Salzburg (SCI-TReCS), Paracelsus Medical University (PMU), Salzburg, Austria; HSCI iPS Core Facility, Harvard University, Cambridge, USA; Department of Transfusion Medicine and SCI-TReCS, PMU, Salzburg, Austria; BCRT & Institute of Medical Immunology, Charité - Univeritätsmedizin Berlin, Germany; NIAID/NIH, Laboratory of Clinical Immunology & Microbiology, Bethesda, MD

**Keywords:** Stromal cells, iPSCs, immunomodulation, immunodeficiency, SCID, CRISPR/Cas9

## Abstract

Stromal cells contribute to organ integrity as fibroblasts and to vascular stability as pericytes, in addition to their enigmatic niche function in many tissues. Their inherent immunomodulatory capacity attracted particular attention, initiating numerous clinical trials, particularly testing trophic regeneration and immunomodulation. Key stromal immune functions are still enigmatic.

Here we show that dedicator of cytokinesis (DOCK-2) previously described for causing immune cell dysfunction plays a role in extra-hematopoietic immunity by regulating stromal/fibroblast immunomodulatory function. We used three independent strategies including iPSC-derived mesodermal stromal cell (MSC) lineage maturation, severe combined immunodeficiency (SCID) patient-derived cells and CRISPR/Cas9 knockout to support our findings.

Human induced pluripotent stem cells (iPSCs) were generated from healthy bone marrow and umbilical cord blood-derived fibroblasts by Sendai virus-mediated transient expression of Yamanaka factors. After mesoderm induction, stromal differentiation was induced by platelet-derived growth factors under animal serum-free conditions. Under feeder-free defined conditions, iPSCs differentiated into expandable and cryo-preservable CD73^+^/CD105^+^/Tra-1-81^−^ early iPS-MSCs lacking immunosuppressive potential. Successive maturation was required for reaching the canonical MSC phenotype and immunomodulatory competence over time, while maintaining clonogenicity, comparable to parental MSCs. Sequential RNAseq revealed acquisition of a spectrum of immune-related genes significantly expressed in mature iPS-MSCs and resembling parental MSC’s immune gene expression. The DOCK-2 gene attracted our attention because mutations can cause SCID. Interestingly, SCID patient-derived fibroblast lines harboring bi-allelic DOCK-2 mutations showed significantly reduced immunomodulatory capacity compared to non-mutated control fibroblasts. CRISPR/Cas9-mediated DOCK-2 knockout in healthy iPSCs resulted in iPS-MSCs that also displayed reduced immunomodulatory capacity, thus confirming a role of DOCK-2 in stromal immune function. At a mechanistic level, DOCK-2 deficiency resulted in disturbed subcellular localization of CDC42.

This provides first evidence for an extra-hematopoietic immunomodulatory role of DOCK-2 in stromal cells, previously considered restricted to hampering immune cell migration/function resulting in SCID. We may speculate that some of the signs and symptoms of persisting immune disease after successful hematopoietic stem cell transplantation in SCID patients could at least in part be due to mutations, like DOCK-2^−/−^, permissive outside the hematopoietic immune system, as evidenced also by the increased virus infection susceptibility of DOCK-2 deficient fibroblasts.

## INTRODUCTION

- Immunity outside immune system / stromal niche functions in immune response modulation
- MSCs are immune modulatory, exact mechanism unknown (incl. anti-septic, hematopoietic niche). Which mechanisms are known? What is still unclear? Donor variation? Tissue of origin variation / exact mechanisms of immune modulation *in vivo* not clear
- Role in maintaining immune homeostasis as niche / maintaining/expanding Tregs,
- Previous iPS-MSCs were mostly lacking or “not testing” immunomodulation → Table + references
- Why iPS differentiation to iPS-MSCs and comparing to parental MSCs? Reprogramming “erases” methylation pattern etc….age, environmental influences, organ specificity?, exposure to LPS, rejuvenation of cells by reprogramming; by comparing MSCs to iPS and iPS-MSCs avoiding donor variation effects
- Add. benefit = comparability of knockouts
- Any role for SCID – discuss post-HSCT outcomes / protracted IVIG need &/or ekcema ?

## RESULTS

To better understand stromal cell immune function, we first devised an ontogeny-inspired model system. Human primary MSCs from bone marrow (BM) and umbilical cord blood (UCB) were selected as starting cells in order to study two different tissues of origin and additionally an adult and a juvenile source of MSCs. Both MSC strains were confirmed previously to display a dose dependent capacity to inhibit T cell proliferation as a surrogate of their immunomodulatory function (*Ref. pubmed/26620155*). These primary MSCs were reprogrammed into iPSCs using a non-integrative Sendai virus protocol leading to the temporary expression of the Yamanaka factors Oct4, Sox2, KLF4 and c-Myc. The MSC-derived iPSCs were induced into mesoderm and thereafter subjected to stepwise differentiation along mesodermal stromal/fibroblast lineage over four to eight passages (p4-8), to study their immune-related phenotype and function in comparison to their parental MSCs (**Fig.1a I**). Single cell immune-phenotyping by multiparameter flow cytometry revealed homogenous loss of pluripotency markers paralleled by a consecutive acquisition of MSC marker expression in the course of differentiation of iPSCs into iPS-MSCs (**Fig.1b, Fig.S1 and S2**).

**Figure 1:**
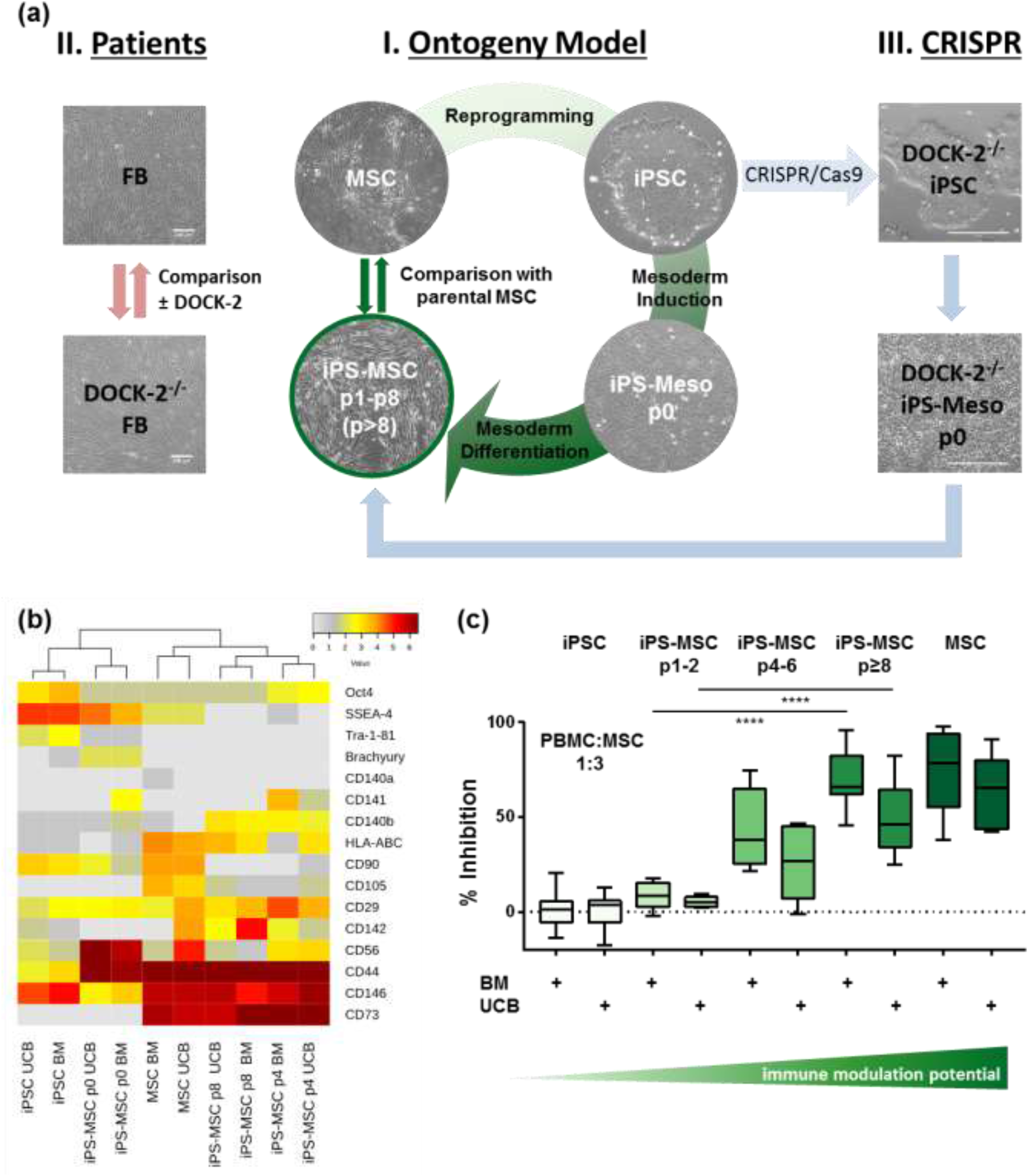
Human iPS-derived MSCs undergo a stepwise maturation leading to a reacquisition of immune modulatory potential. **(a)** Experimental setup: **I.** MSCs derived from bone marrow (BM) and umbilical cord blood (UCB) were reprogrammed into iPSCs before mesoderm induction and stepwise maturation to mesodermal stromal cells (iPS-MSCs) to enable sequential gene expression profiling. **(b).** Cluster analysis of single cell-based flow cytometric marker expression profile during differentiation from iPSCs to iPS-MSCs compared to parental MSCs. Mean values of delta MFI (specific antibody staining minus isotype control) from passage 0 to passage 8 are shown. **(c)** The immune modulatory potential of iPS-MSCs inhibiting T-cell mitogenesis increased with the degree of differentiation from early to late differentiation passages. BM- and UCB-derived progeny showed a significant anti-proliferative effect from passages four and beyond comparable to their parental primary MSCs. (BM n = 6-10; UCB n = 5-8; PBMC:MSC ratio 1:3; one way ANOVA including Tukey’s multiple comparison test ****p < 0.0001). Percentage of inhibition after normalizing the mean values of T cell proliferation of single assays are shown. Experimental setup: **II.** DOCK-2 deficient fibroblast (FB) lines were compared to healthy control FBs. **III.** Healthy iPSCs from setup I were used for CRISPR/Cas9-mediated bi-allelic DOCK-2 knockout and subsequent DOCK-^2−/−^ iPS-MSC propagation and additional comparison.

Most iPSC-derived MSCs described in previous studies lacked the capacity to temporarily inhibit lymphocyte proliferation (**Table S1**). We therefore compared mesoderm-derived iPS-MSCs at early vs. later passages directly to their parental MSCs and MSC-derived iPSCs and observed a gradual significant increase of MSC-dependent inhibition of T cell proliferation over time (**Fig.1c**).

Human iPSCs derived from BM-MSCs and UCB-MSCs showed a normal karyotype after Sendai virus elimination and differentiated into endodermal, ectodermal and mesodermal tissue forming a teratoma in immunodeficient mice (**Fig.S4a–c**). Mesoderm induction was performed for four clones of BM- and three clones of UCB-derived iPSC lines and was confirmed by upregulation of CD56 and Brachyury (**Fig.S5+S6**; *Ref. Evseenko PNAS, 2010*). The expression of pluripotency markers Oct4, Tra-1-81 and SSEA-4 decreased during the differentiation while Brachyury and CD56 proteins were most abundant in iPS-MSCs p0 immediately after mesoderm induction. The expression of CD56 was documented after mesodermal induction and was reproducibly and significantly enhanced in iPS-MSCs (**Fig.1b, S1+S5**). The significant upregulation of MSC-related molecules including CD73 (5’-nucleotidase), CD105 (endoglin) and reduction of Tra-1-81 during differentiation (**Fig.S1**) was paralleled by a consecutive acquisition of the typical fibroblastoid morphology comparable to the parental cells (**Fig.S7+S8**). Reproducible gain of CD146 (melanoma cell adhesion molecule, MCAM) (*Ref. Bianco et al, CSC 2012*), CD140b (platelet-derived growth factor receptor beta, PDGFRb; *Ref. Scheding CSC 2015*), CD142 (tissue factor; *Ref. Öller THN 2018*), CD29 (integrin β1), CD44 (phagocytic-glycoprotein 1) and HLA-AB (human leukocyte antigen alleles A, B, representing major histocompatibility complex antigens class I, MHC I) indicated progressive differentiation into iPS-MSCs with increasing passages (shown are p0, p4 and p8; **Fig.1b and S1**). The colony forming capacity of iPS-MSCs was maintained (**Fig.S9**) and trilineage differentiation capacity was reacquired after mesodermal differentiation, although with limited chondrogenic differentiation compared to parental BM-MSCs (*Ref: pubmed/25406351*; **Fig.S10**).

Comprehensive consecutive gene expression profiling of multiple clones by RNA sequencing (RNAseq) comparing key differentiation steps identified 4,820 genes to be significantly regulated during iPS-MSC maturation into a characteristic parental adult stromal cell expression status, thus separating MSCs from iPSCs and early mesoderm (p0; **Fig.2a**). Principle component analysis (PCA) identified a tremendously stable developmental process covering 66% and 22% variance in the two dominant principle component (PC)-forming gene clusters PC1 and PC2, respectively, and including the re-acquisition of the parental gene expression profile (**Fig.2b**). The top 500 regulated genes in the time course of iPS-MSC ontogeny also reflected a developmental path with iPS-MSC re approaching their adult predecessor’s gene expression profile (**Fig.S11b**). Among the 200 most significantly up-regulated genes belonging to the gene ontogeny categories “immune system process” we focused on those 25% of genes related to T cell immunity (**Fig.S11b**) including two striking gene clusters regulating T cell immunology and immune-related G-protein coupled receptor (GPCR) signalling (**Fig.2c**). DOCK-2 attracted particular attention because mutations of this guanine exchange factor (GEF), commonly involved in Rho kinase regulation, were recently discovered to result in a “combined immunodeficiency with pleiotropic defects of hematopoietic and nonhematopoietic immunity” (*Ref: Dobbs/Notarangelo NEJM 2015, J Clin Immunol 2019*). *This may point to* a missing link between immune cell dysfunction and the still enigmatic immune response modulation capacity of the stromal/fibroblastoid niche cells (*Ref: Ankrum/Karp, nbt 2015; PMC4320647/*). Additional regulation of the Ras-related C3 botulinum toxin substrate 2 (RAC2) gene and several other immunologically relevant G-protein signalling molecules during MSC ontogeny from pluripotent stem cells (**Fig.2c**) supported the hypothesis that the otherwise stromal niche-forming “MSCs” also play a role in immunity.

**Figure 2:**
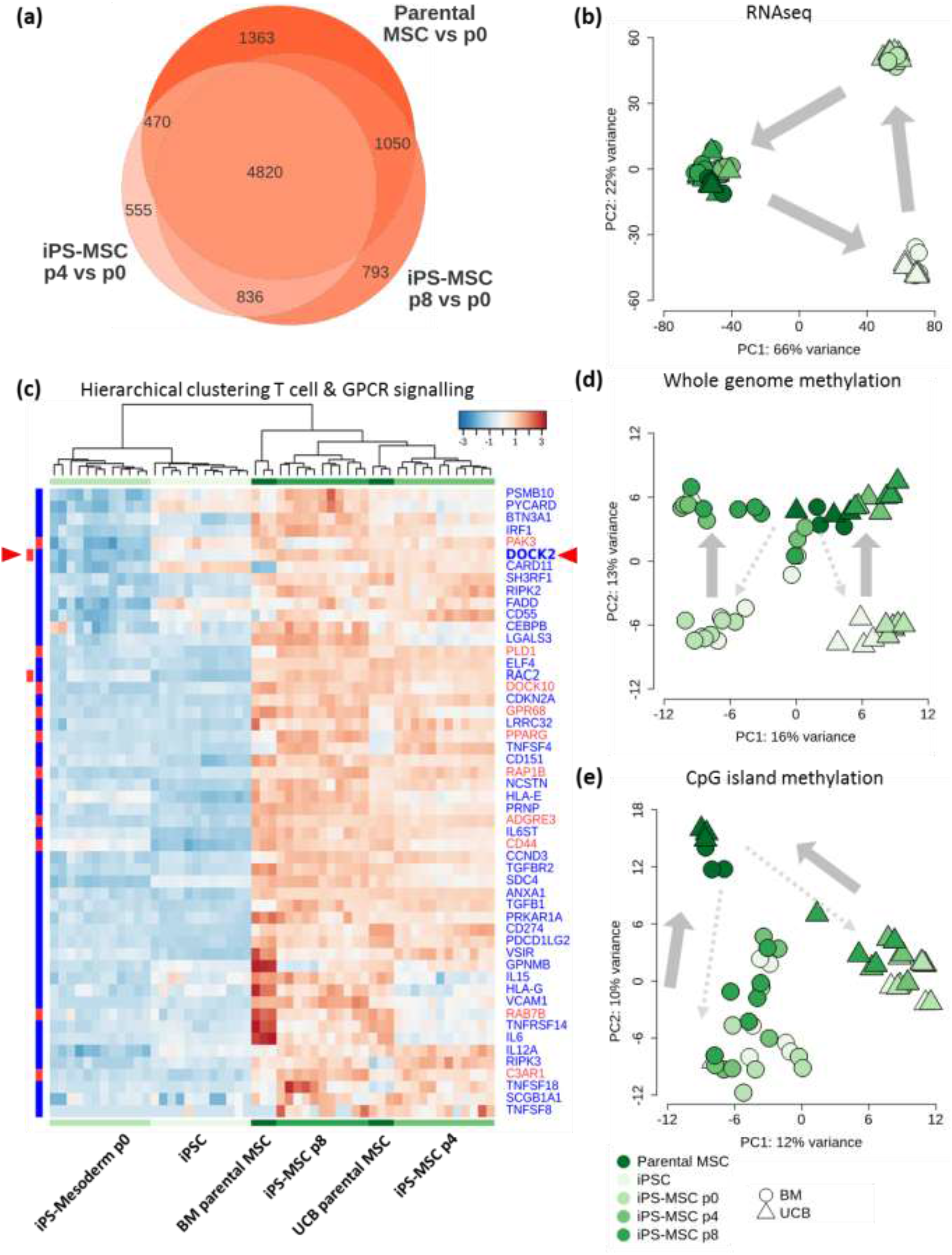
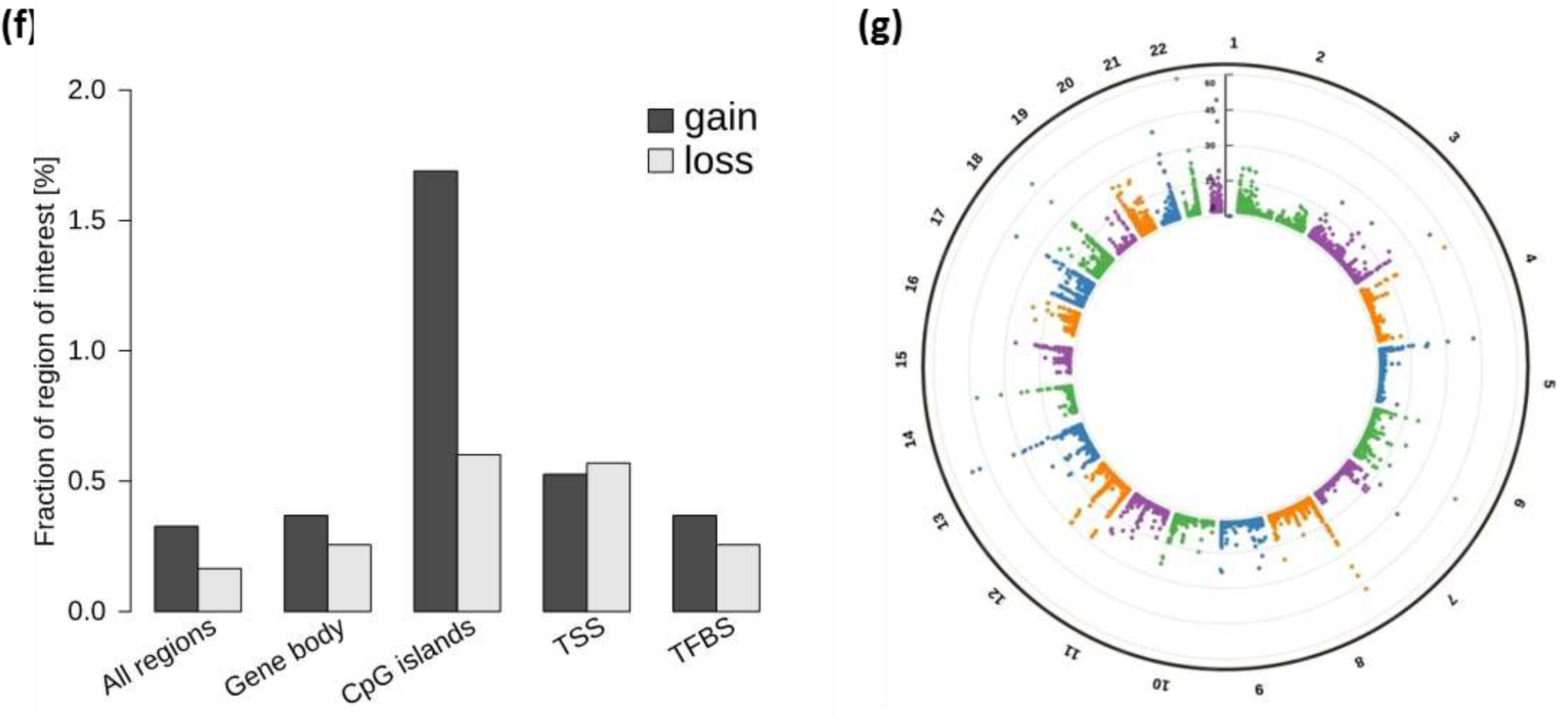
Sequential gene expression and epigenomic analysis during iPS-MSC ontogeny. **(a)** Venn diagram summarizing the number of significantly differentially regulated genes (>2-fold) in parental MSCs vs. iPS-MSCs p0 and iPS-MSCs p4 vs. p0 and p8 vs. p0 (adjusted p-value < 0.05). 4820 genes (25% of the 19,287 detectable gene IDs) of the differentially expressed genes were commonly found in all three comparisons. **(b)** Hierarchical clustering heat map of genes linked with <u>T-cell proliferation</u> (**blue**) or G-protein signaling related (**red**) and found significantly upregulated in iPS-MSCs p8 vs. p0 iPS-mesoderm and also represented in parental MSCs with two genes localized in both categories (DOCK2 and RAC2). (**c**) Principal component (PC) analysis at the transcriptome and (**d**) whole methylome levels and (**e**) CpG island methylation sub-analysis was used to explore the relationship between different samples. Mature iPS-MSCs group together with their parental BM and UCB-derived MSCs in both RNAseq and genome methylation principle component analysis (PCA). The different donors and clones group together at methylation level. *Most variation is explained by the grade of differentiation*. The grey arrows track the differentiation path. P values were adjusted using Benjamini-Hochberg correction. (RNAseq: iPSCs n = 12; p0 iPS-mesoderm n = 12; p4 iPS-MSCs n = 12; p8 iPS-MSCs n = 11; parental MSCs n = 6; Methylseq: iPSCs n = 11; p0 iPS-MSCs n = 12; p4 iPS-MSCs n = 9; p8 iPS-MSCs n = 10; parental MSCs n = 6). (**f**) Feature enrichment showing the percentage of regions of interest (all regions, gene body, CpG islands (Cpg), transcription starting sites (TSS) and transcription factor binding sites (TFBS)) gained or lost between iPS-MSCs p8 and p0. (**g**) Circular Manhattan blot showing the differentially methylated regions between iPS-MSCs p8 and p0 on the whole genome with chromosome numbers on the external ring. Each dot corresponds to a differentially methylated region (500 base pair regions). The height of the dot correlates with the level of significance (log 10 adjusted p-value, Benjamini Hochberg correction).

<u>As a second experimental strategy</u>, we therefore tested DOCK-2-deficient fibroblast lines from two SCID patients that were previously described to display enhanced virus replication and viral-induced cell death after experimental encephalomyocarditis virus infection (rescued by alpha interferon treatment or lentiviral DOCK-2 transduction) for their capacity to inhibit/modulate T cell proliferation (**Fig.1a II and Fig.S12**). The patient fibroblasts showed a dose-dependent significantly reduced inhibition of T cell mitogenesis (**Fig.3a**) and allogeneic mixed lymphocyte reaction (**Fig.3b**). Direct comparison of these results to the iPS-MSC’s and their parental MSC’s immunomodulatory capacity (**Fig.1c**) was hampered naturally by donor variation as well as organ/tissue source and age variability of the different stromal cells (*Ref. Reinisch/Irv Weissman/Strunk Blood 2015*).

**Figure 3:**
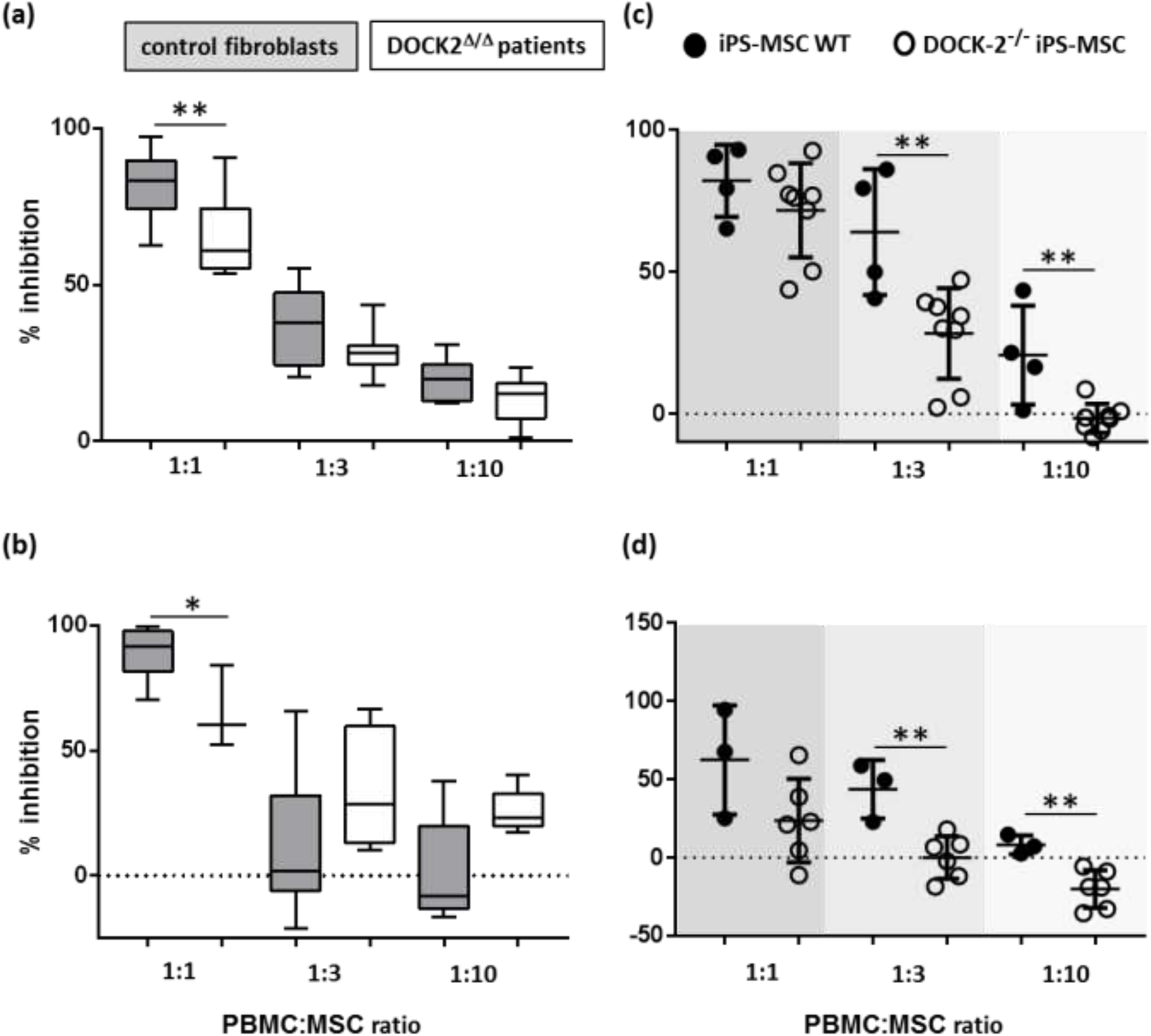
Fibroblasts of DOCK2 deficient patients and CRISPR/Cas9-created DOCK-2^−/−^ iPS-MSCs showed impaired immune modulatory potential. (**a**) Fibroblasts were co-cultured with CFSE-labelled PBMC and T cell mitogenesis was analysed at day 4. Pooled data of the two DOCK2 patient’s fibroblasts from repetitive experiments and four different adult donor’s fibroblasts (n = 9-11, **p < 0.01, unpaired two-tailed T test 1:1 p= 0.0076, 1:10 p=0.029). (**b**) Fibroblasts were co-cultured with PBMCs in an allogeneic mixed leukocyte reaction (MLR) and proliferation was analysed at day 7. Pooled data of the two DOCK2 patients, 5 different adult fibroblast and 2 neonatal fibroblast donors (n = 5-11, *p < 0.05 unpaired two-tailed T test p=0.0149)). (**c**) Wild type (WT) iPS-MSCs and DOCK-2^−/−^-iPS-MSCs inhibiting T cell mitogenesis and (d) MLR in a dose dependent manner. Mean values of T cell proliferation triplicates of single assays and the percentage of inhibition after normalizing the data to (a, c) maximum proliferation after phytohemagglutinin stimulation or (b, d) MLR. Statistics (c) n = 4-8, unpaired two-tailed T test 1:3 p=0.0090; 1:10 p=0.0059); (d) n = 4-6, unpaired T test 1:1 p=0.0398; 1:3 p=0.0016, 1:10 p=0.0029).

<u>A third strategy using CRISPR/Cas9-mediated bi-allelic DOCK-2 knockout</u> was thus used for creating DOCK-2 deficient cell clones directly comparable to their iPS-MSC ‘siblings’ derived from the same initially picked iPSC clone (UCB-iPSC clone ‘C’). This guided the initial discovery of DOCK-2 expression paralleling the acquisition of the MSC’s immunomodulatory capacity during mesodermal stromal ontogeny in this study (**Fig.1a III and Fig.S14**). Direct head-to-head assessment confirmed a dose-dependent significant lack of T cell proliferation inhibition by DOCK-2^−/−^ iPS-MSCs compared to healthy UCB-derived iPS-MSCs (**Fig.3c+d**).

(**Fig.2d-e**) incomplete regeneration of epigenomic identity corresponding to the lack of complete surface marker identity (CD90) that required extended culture, i.e. final complete phenotypic maturation, achieved beyond passage eight (**Fig.S13**).

(**Fig.2f-g**) Pronounced CpG island methylation was observed during iPS-derived mesodermal stromal ontogeny, despite incomplete re-establishment of the parental epigenomic signature; interestingly none of the 53 selected predominantly regulated genes affecting T cell immunity (blue) and GPCR signalling(red; **Fig.2c**) was regulated significantly at the epigenomic level (**Fig.2g-h**), indicating that …

## DISCUSSION

MSC-derived immunomodulation was already observed in multiple clinical clinical trials - but - it is not clear yet if there is a natural role of a stromal immunomodulatory function beyond increased viral susceptibility in SCID (*Ref: Dobbs/Notarangelo NEJM 2015*)?

…

**Figure 4:**
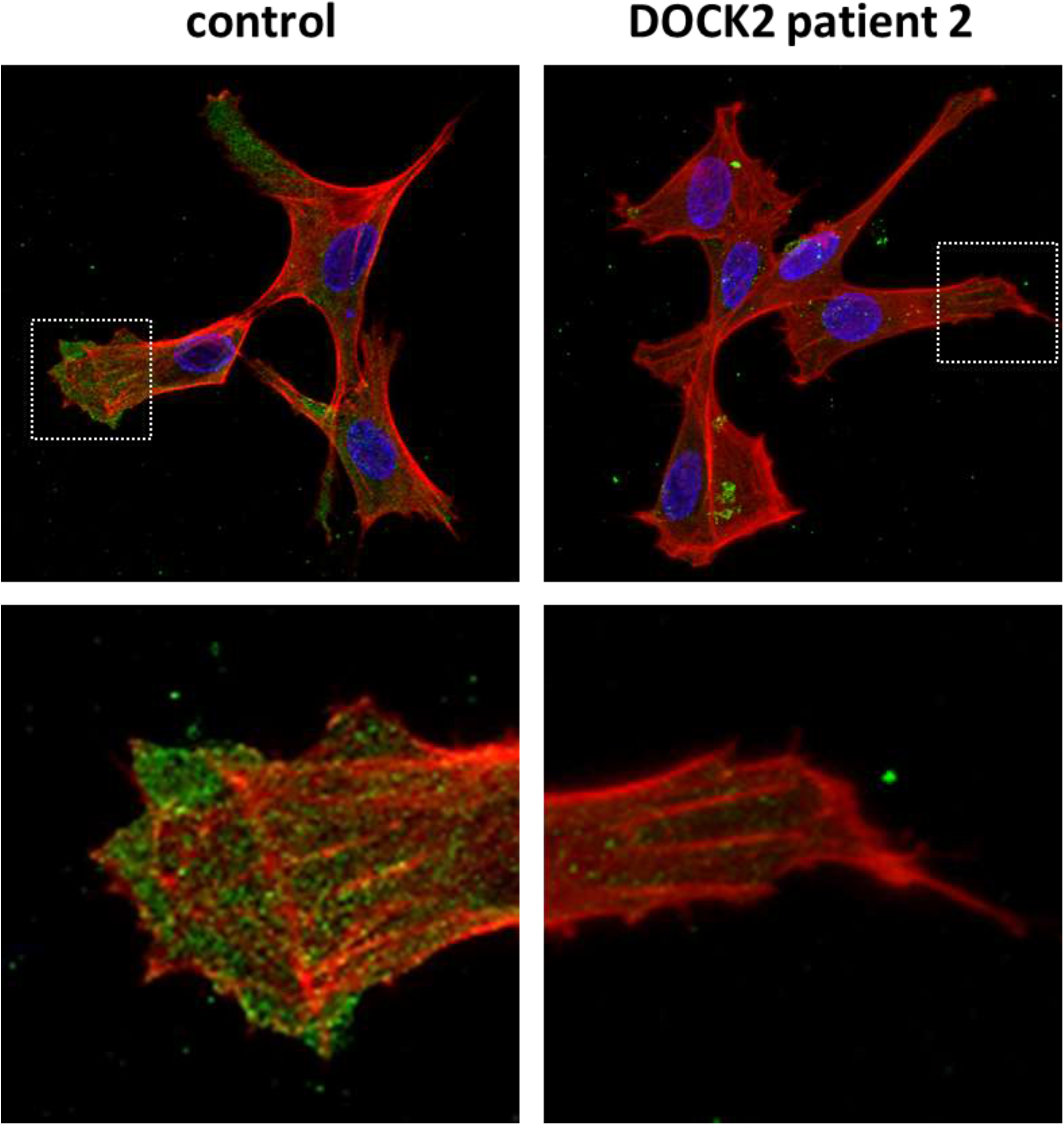
Subcellular localization of CDC42 / F-Actin in control fibroblasts and DOCK-2 deficient patient fibroblasts. Confocal microscopy of skin fibroblasts (control) and patient fibroblast line cells stained for CDC42 (green) and Phalloidin (red); nuclear counterstain DAPI (blue) - add. controls incl. control iPS-MSCs p8 and CRISPR^−/−^ iPS-MSCs p8 still missing = work in progress …

## MATERIALS & METHODS

### Cell isolation, reprogramming and culture

Approval was obtained for human cell and tissue sample collection and genetic reprogramming from the Institutional Review Board (protocols 19–252, 18–243, 21–060, 19–284 and 415-E/1776/4-2014, Ethics Committee of the province of Salzburg). Adult samples were collected in accordance with the Declaration of Helsinki after written informed consent from healthy volunteers. Umbilical cord blood (UCB) samples were collected after written informed consent by the mother-to-be obtained prior to delivery of full-term pregnancies. MSCs from bone marrow (BM) and UCB were isolated and expanded under animal serum-free conditions using pooled human platelet lysate (pHPL) replacing fetal bovine serum and their purity, identity, and viability was characterized by flow cytometry as previously described [https://www.ncbi.nlm.nih.gov/pubmed/17655588 https://www.ncbi.nlm.nih.gov/pubmed/19321860, https://www.ncbi.nlm.nih.gov/pubmed/18620484, https://www.ncbi.nlm.nih.gov/pubmed/19816400]. Clonogenicity and differentiation capacity was assessed as previously described [ https://www.ncbi.nlm.nih.gov/pubmed/17635045]. PBMC were isolated by density centrifugation from random donor buffy coats as described [https://www.ncbi.nlm.nih.gov/pubmed/26620155].

Induced pluripotent stem cells (iPSCs) were reprogrammed from primary MSCs (derived from BM or UCB) by non-integrative Sendai Viral vector kit (CytoTune™-iPS Sendai Reprogramming Kit encoding for Oct4, Sox2, KLF4 and c-Myc, Life Technologies Cat.No.A1378001) *by the Harvard Stem Cell Institute (HSCI) iPS Core Facility (Cambridge, MA, USA)*. *The reprogramming protocol was described by Fusaki and colleagues in 2009* [https://www.ncbi.nlm.nih.gov/pubmed/19838014] *and adapted by the HSCI*. Human iPSCs were characterized by cytogenetic analysis of at least twenty G-banded metaphase cells per clone and teratoma formation.

Human iPSCs were initially transferred from mouse embryonic feeder layers to feeder-free conditions and thereafter maintained on a selected batch of Matrigel® (Corning) in mTeSR™1 (STEMCELL Technologies) medium. When reaching a cell density appropriate for splitting, iPSCs were harvested using Gentle Cell Dissociation Reagent (STEMCELL Technologies). Viable cells were counted and seeded at a density of 5 × 10^4^ cells per cm^2^ on Matrigel® in mTeSR™1 containing 10 μM Y-27632 ROCK pathway inhibitor (Selleckchem). After 24 hours, the medium was changed to STEMdiff™ Mesoderm Induction Medium (STEMCELL Technologies, *defined, xeno-free medium*) and the medium was replaced daily for the next consecutive 4 days. On day 5, cells were harvested using TrpLE™ (Gibco) and the expression of early mesoderm markers Brachyury and CD56 was determined by flow cytometric analysis. Cells were seeded in EGM™-2 containing hydrocortisone, hFGF-B, VEGF, R3-IGF, Ascorbic Acid, hEGF (Lonza), preservative-free heparin 10 IU/mL (Biochrom), dipeptiven (1.2 mg/ml, L-glutamin – L-arginin dipeptide, clinical grade: Fresenius Kabi), 10% pHPL and 10 μM Y-27632 at a cell density of 27,000 cells/cm^2^ without Matrigel® coating and cultured in humidified incubators (Binder CB210) at 37°C and 5% CO2 in ambient air. Until passage five, iPS-MSCs were passaged at a density of 27,000 cells/cm^2^ twice per week with addition of Y-27632. For microscopical documentation EVOS XL (Thermo Fhisher Scientific) was used. Differentiations were performed with iPS reprogrammed from MSCs derived from bone marrow (4 clones) or umbilical cord blood (3 clones).

### Flow cytometry immune phenotyping and T cell proliferation assay

Immune phenotyping of MSCs, iPSCs and iPS-MSCs was performed using a BD LSRFortessa™ (Becton Dickinson) and the following antibodies with their corresponding isotype controls: CD19-BUV395, CD29-APC, CD44-PE, CD45-APC, CD73-PE, CD90-BUV395, CD140a-BV421, CD140b-BV421, SSEA-4-PE, Tra-1-81-Alexa Fluor 647, HLA-ABC-BUV395 (BD), CD14-PE, CD31-eF450, CD34-PE-Cy7, CD56-PE, CD105-eF450, HLA-DR-eF450 (eBioscience), CD141-APC, CD146-PE-Vio770 (Milteny), Brachyury-APC (R&D Systems) and Oct4-PE (Biolegend).

Dead cells were excluded based on FVD-eFluor™520 (eBioscience™) staining. For intracellular staining Fix & Perm Solution (eBioscience™) was used according to the manufacturer’s protocol. Results were analyzed using Kaluza Analysis Software (Beckman Coulter).

Immunomodulatory potency of MSCs, iPSCs and iPS-MSCs was determined as described [https://www.ncbi.nlm.nih.gov/pubmed/?term=Ketterl=2015]. Briefly, peripheral blood mononuclear cells (PBMC) from ten random donors were pooled, stained with carboxyfluorescein succinimidyl ester (CFSE, 2 μM, 15 minutes, 37 °C; Sigma) and cryopreserved in liquid nitrogen in order to have reference responders in multiple subsequent experiments. For immune modulation assays; 3 × 10^5^ CFSE pre-labeled PBMC resuspended in RPMI-1640 supplemented with 10% pHPL (for all assays performed with iPS-MSCs) or AB serum (for all assays performed with DOCK2 knockout cells), 2 IU/mL preservative-free heparin (Biochrom), 2 mM L-glutamine (Gibco), 10 mM HEPES (Gibco), 100 IU/mL penicillin and 100 μg/mL streptomycin (Sigma) were plated per well in triplicate in 96-well flat-bottomed plates (Corning). T cell proliferation in four-day mitogenesis assays was induced by 5 μg/ml phytohemagglutinin (PHA; Sigma). Allogeneic mixed leukocyte reactions due to the pooling of 10 independent PBMC donor-derived cells were measured at day seven. PBMC were cultured with or without graded numbers of MSCs (250 μL total volume per well) in threefold serial dilution as indicated in the results section. All cultures were performed in humidified ambient air incubators (Binder CB210) at 37°C and 5% CO_2_ in ambient air. Proliferation of viable CD3^+^ cells was analyzed using using a Gallios 10-color flow cytometer and the Kaluza G1.0 software (both Coulter). Viable 7-aminoactinomycin-D-excluding (7-AAD; BD Pharmingen) CD3-APC^+^ (eBioscience) T cells were analyzed.

### RNASeq, MethylSeq and bioinformatics

Libraries were sequenced on a Hiseq 1500 (Illumina) with 50 bp single-end reads. Quality control was conducted using FASTQC. Trimmomatic was used to remove residual adapter sequences and remove low quality reads. Reads were then mapped to the Ensembl GRCh38 human genome using Bowtie 2 (ask Karsten for confirmation). The number of mapped reads/gene (counts) was then calculated using HTseq. Genes were annotated using the Ensembl version 97. Expression values of protein coding genes were normalized using Deseq2 package. In order to see, how samples cluster together, a principal component analysis (PCA) and hierarchical clustering analysis using Euclidean distance were conducted on the whole normalized dataset. Differential expression analysis between the different groups was conducted using Deseq2. Genes with an adjusted p-value < 0.05 (Benjamini Y, Hochberg Y (1995) Controlling the false discovery rate: a practical and powerful approach to multiple hypothesis testing. J R Stat Soc B 57:289–300 https://www.jstor.org/stable/2346101?seq=1/subjects) were considered significantly differentially transcribed. Enrichment analysis (Go term enrichment, Gene set enrichment analysis, Kegg pathways and Panther pathways) were conducted using “clusterProfiler” package in R. Benjamini and Hochberg multiple testing correction was used to adjust raw p-values for multiple testing (adj. p-value < 0.05 were considered significant).

### Scratch assay

For wound repair studies, fibroblasts were seeded at cell density of 20.000 cells per well in 24 well plates and cultured until confluence (around 24 hours). Cells were serum starved overnight in 0.2% HPLS. After standardized scratch of the confluent layer with a 200 μl pipette tip, medium was refreshed and cultures were introduced into a Okolab incubator system surrounding a Nicon Eclipse Ti. Cell movement was monitored by acquiring video sequences using NIS-Elements software covering a time period of 12 hours. The area of wound repair was determined using TScratch software (Gebäck Biotechniques 2009).

### Immunofluorescence

Cells were seeded on collagen-coated glass coverslips. Cells were fixed with 4% PFA for 15 minutes at room temperature, then washed in PBS and permeabilized for 10 min in 0.1% TritonX100/PBS. After 30 min blocking in 10% FBS/Dako wash buffer, anti-CDC42 antibody (Abcam, ab187643) or the corresponding isotype control (Abcam ab172730) were applied 1:500 and incubated overnight at 4°C. After washing in Dako wash buffer, the Phalloidin Alexa Flour 568 (1:40, Invitrogen) and the goat-anti-rabbit-Alexa Fluor 488 secondary antibody (1:500, Invitrogen) were applied in 10% FBS/Dako wash buffer for 1h at RT. Finally, the coverslips were washed in Dako buffer and mounted in RotiMount FluorCare (Roth) including Dapi on glass slides. Confocal pictures were taken using a Zeiss LSM 710 and ZEN software.

### DOCK2 knockout

For introduction of a knockout of the DOCK2 gene (NCBI gene ID 1794) the web based tool CRISPOR (http://crispor.tefor.net/) was used to identify possible guide sequences that target exon 37 of the DOCK2 gene. Two guide sequences, for which CRISPOR calculated low off-target binding sites and high cutting efficiencies, were selected for experimental evaluation (see **table S2** for sequences). Subsequently, the selected guides were tested for their cutting efficiency and indel formation frequency by transfection of the hiPSC, sequencing of the target site and analysis using the TIDE algorithm (https://tide.deskgen.com/). Methodical details are described below. Application of guide-2 resulted in the higher cutting and indel formation frequency (69,4%) and was therefore used for the generation of the knockout clones.

For editing the Alt-R CRISPR-Cas9 System (Integrated DNA technologies) was used. This included the following reagents: Alt-R® S.p. Cas9 Nuclease V3 (Cat. 1081058), Alt-R® CRISPR-Cas9 tracrRNA (Cat. 1073191) and the crRNA was (crRNA: GAAGATCGCGGAGTTTGTAC). The gRNA duplex was generated by formation of *crRNA:tracrRNA duplex.* Briefly 5μl of the specific crRNA (100μM) and 5μl of tracrRNA (100μM) were mixed and incubated for 5 minutes at 95°C followed by down cooling to RT for 15 min. The RNP-complex was formed by mixing 2μl of gRNA duplex and 2μl of Alt-R® S.p. Cas9 Nuclease V3 (61μM) and incubation for 35 minutes at RT.

Transfections were carried out using the P3 Primary Cell 4D-Nucleofector X Kit (Lonza, Cat. V4XP-3024) with program CM150 in the Lonza Nucleofector™ device (Core Unit and X Unit). Briefly the hiPSC were harvested by incubation with TrypLE (Thermo, Cat.12563-029) to yield a single cell suspension. A total cell number of 1,3×10E6 cells were resuspended in 100μl of transfection buffer mix of the kit, 4 μl of the RNP-complex was added and transfected usine program CM150 of the nucleofection device (Lonza). After transfection cells were seeded in StemFlex™ medium (Thermo Fisher, Cat. A3349401) supplemented with 10% CloneR (Stem Cell Technologies Cat. 05888) into one well of a Geltrex (Thermo Fisher Cat. A1413202) coated 6 well plate. Culture medium was changed daily with addition of CloneR for the first three days after transfection. On day four after transfection, the medium was switched to E8 medium (Chen G, et al. 2011). To isolate clones, cells were passaged using TrypLE and seeded as single cells in Geltrex coated 6 well at a density of 50 - 300 cells/well. Individual hiPSC colonies from these cultures were picked into the wells of a Geltrex coated 24 well plate using a pipet tip. After four days of culture cells were harvested from each well by 0,5 mM EDTA treatment. Half of the individual cell suspensions were frozen using Bambanker freezing medium (Nippon Genetics Europe, Cat. BB01-NP) and stored in liquid nitrogen and the other half was used to amplify the targeted genomic region using the Phire Animal Tissue Direct PCR Kit (Thermo Fisher, Cat. F-140WH) regarding the manufacturer’s instructions (**Table S2**) for primer sequences).

PCR products were analysed using 1,5% agarose gel electrophoresis to confirm amplification and allow subsequent clean up using the NucleoSpin Gel and PCR Clean-up Kit (Macherey Nagel, Cat. 740609.250). Samples were subjected to Sanger Sequencing (Microsynth Seqlab) and results were analysed using Snapgene software (GSL Biotech LLC). Three clones were identified to carry a frameshift mutation resulting in a premature stop codon.

**Table S1:**
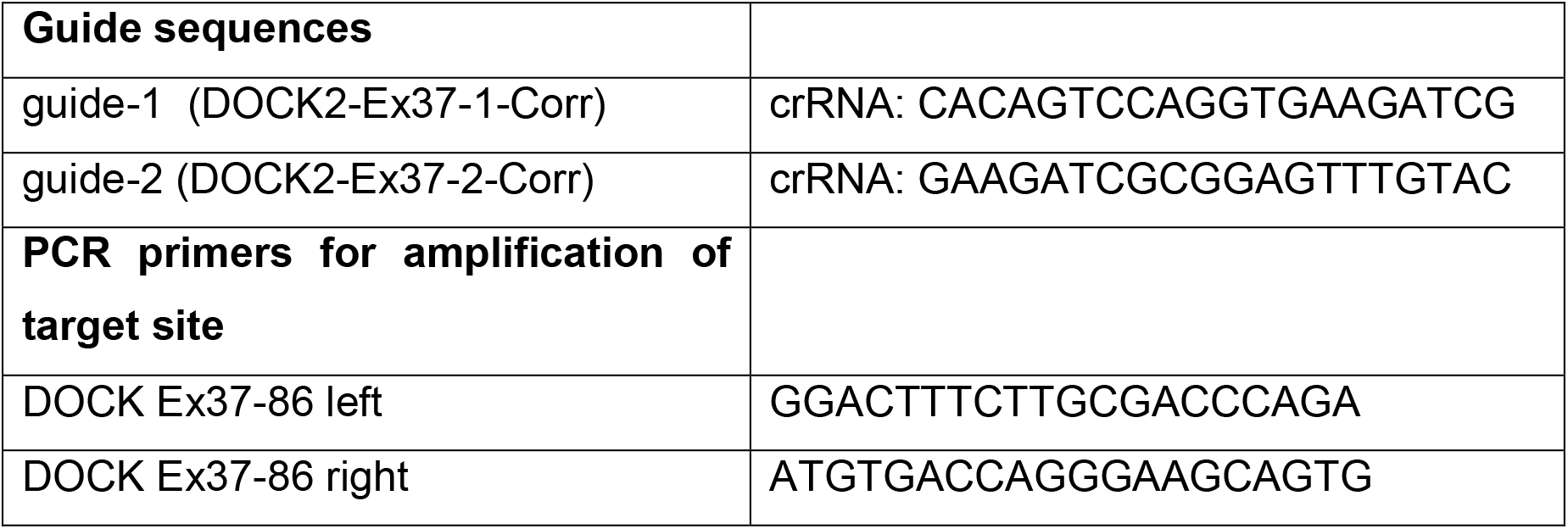

### Statistics

Statistical analysis of the results was performed using One-Way ANOVA analysis of variance with a confidence interval of 95% and corrected for multiple comparisons using the Holm Sidak algorithm in GraphPad Prism version 7.03. Proteomic results were analyzed using R. A p value of <0.05 was defined as significant.

4,820 genes to be significantly regulated during iPS-MSC ontogeny:

- Three comparisons

Parental MSC vs P0 & P8 vs P0 & P4 vs P0 UP 2115
Parental MSC vs P0 & P8 vs P0 & P4 vs P0 DOWN 2543
- Two comparisons

P8 vs P0 & P4 vs P0 UP 2575
P8 vs P0 & P4 vs P0 DOWN 2988
Parental MSC vs P0 & P4 vs P0 UP 2310
Parental MSC vs P0 & P4 vs P0 DOWN 2731
Parental MSC vs P0 & P4 vs P0 UP 2536
Parental MSC vs P0 & P8 vs P0 DOWN 3150
- One comparisons

Parental MSC vs P0 UP 3450
Parental MSC vs P0 DOWN 4253

## Acknowledgements

Luigi D Notarangelo and Kerry Dobbs (NIH) generously provided SCID patientderived fibroblast lines. We are grateful to Nina Ketterl, Anna Hoog and Anna M Raninger for excellent technical assistance.

## DATA SUPPLEMENT

**Figure S1:**
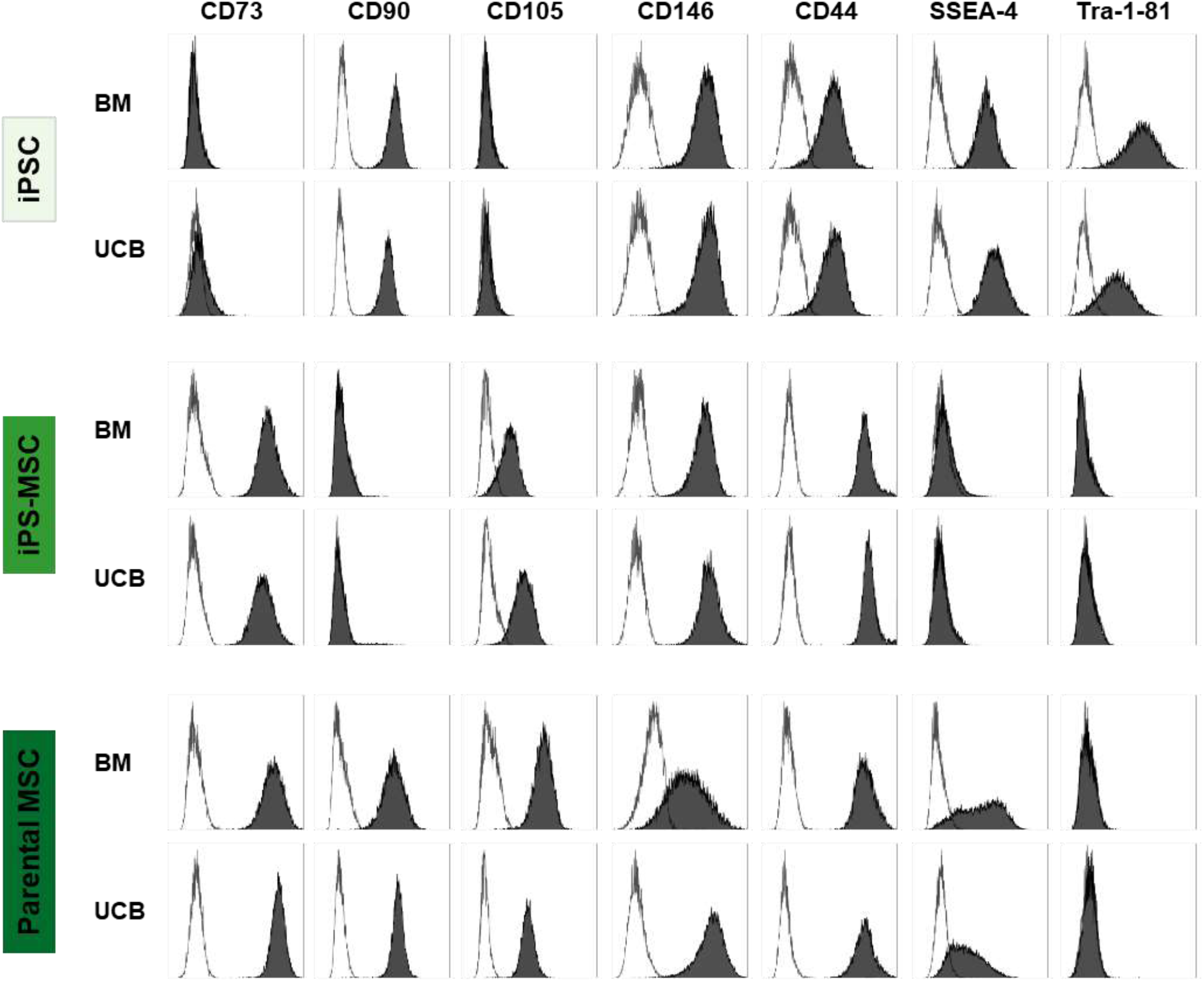
Marker expression profile in the course of differentiation of iPSCs into iPS-MSCs. Representative histograms of CD73 expression in iPSCs and iPS-MSCs p8 compared to parental MSCs. Histograms show fluorescent cell surface staining of anti-CD73 monoclonal antibody conjugated to a fluorophore (gray shading) and its corresponding isotype control (no shading). CD73 was clearly re-upregulated in iPS-MSCs p8 after differentiation and was comparable to parental MSCs in BM and UCB-derived cells. Histograms show the populations (% gated) obtained following the hierarchical gating strategy: size and granularity, doublet exclusion, live cell population and corresponding marker.

**Figure S2:**
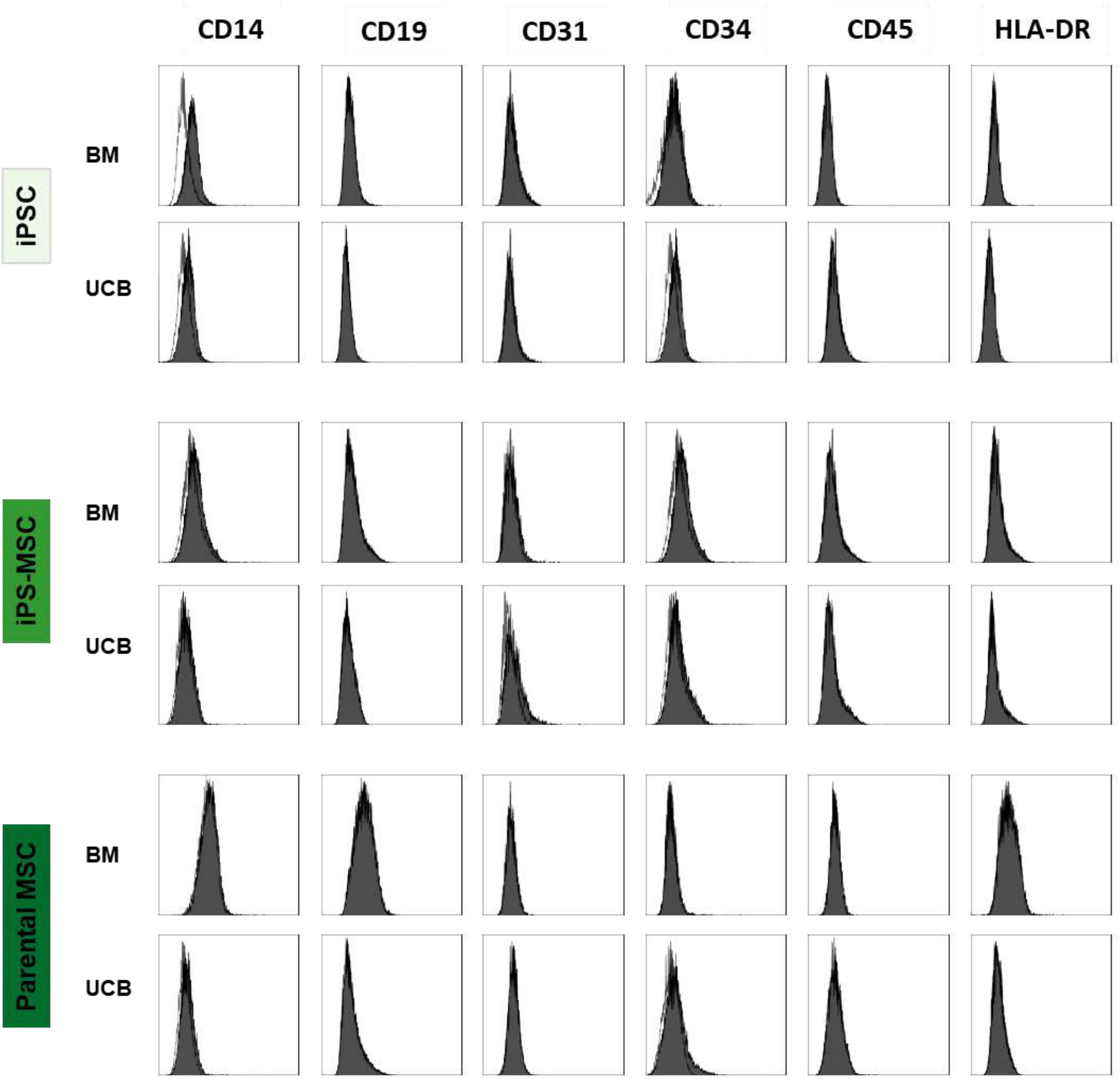
Lack of markers of the hematopoietic lineage in MSCs, iPS and iPS-MSCs. Shown are representative histograms of fluorescent surface staining of anti-CD14, anti-CD19, anti-CD31, anti-CD34, anti-CD45 and anti-HLA-DR monoclonal antibodies conjugated to fluorophores (gray shading) and their corresponding isotype control (no shading) of MSCs, iPS and differentiated iPS-MSCs p≥8 derived from BM or UCB. MSCs, iPSCs and iPS-MSCs do not show expression of PBMC or endothelial markers.

**Figure S3:**
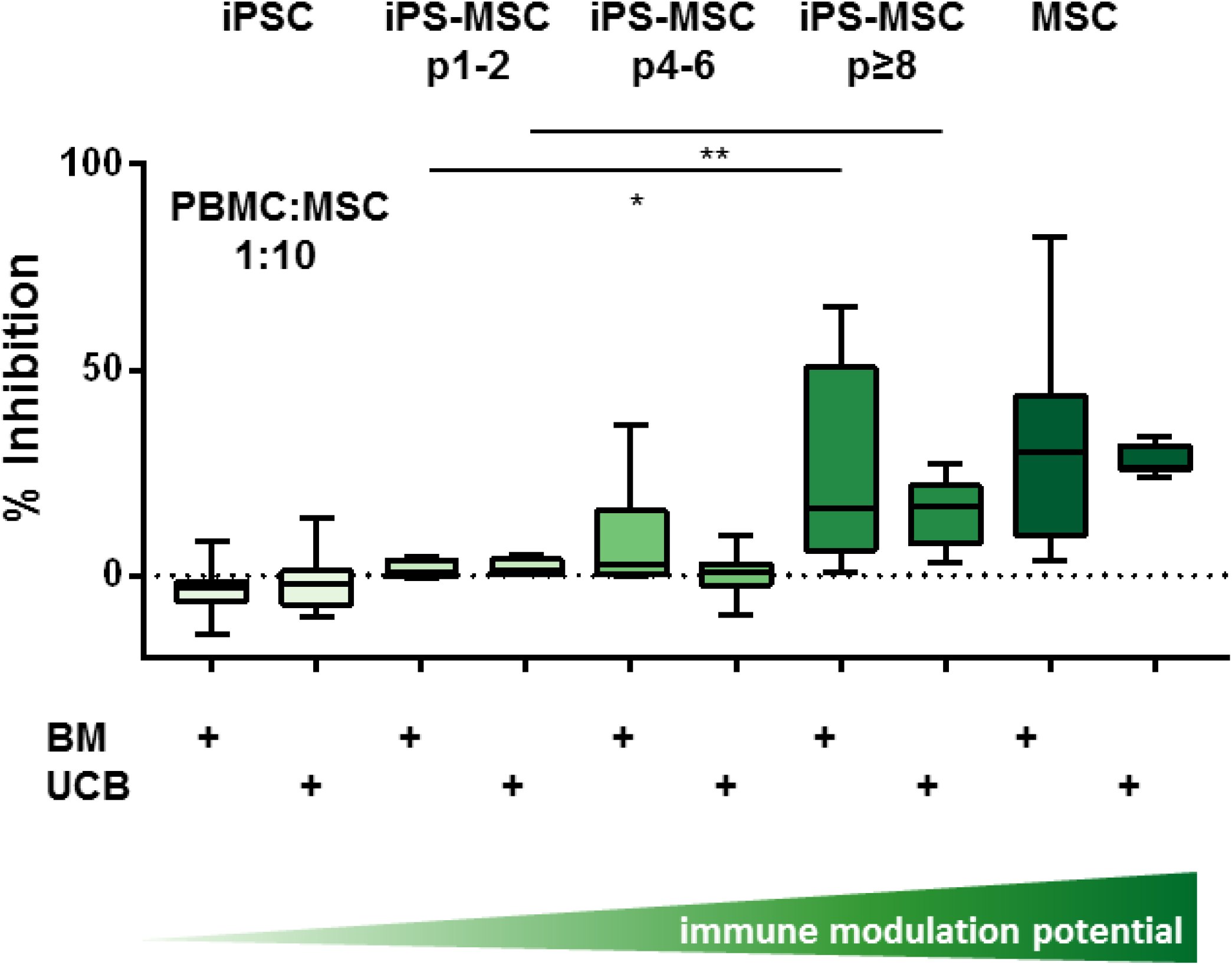
Dose dependent inhibition of T cell mitogenesis by iPS-MSCS (1:10 ratio). Corresponding to Fig.1c …(unpaired T test BM p=0.0433; UCB p=0.0087)

**Table S1:**
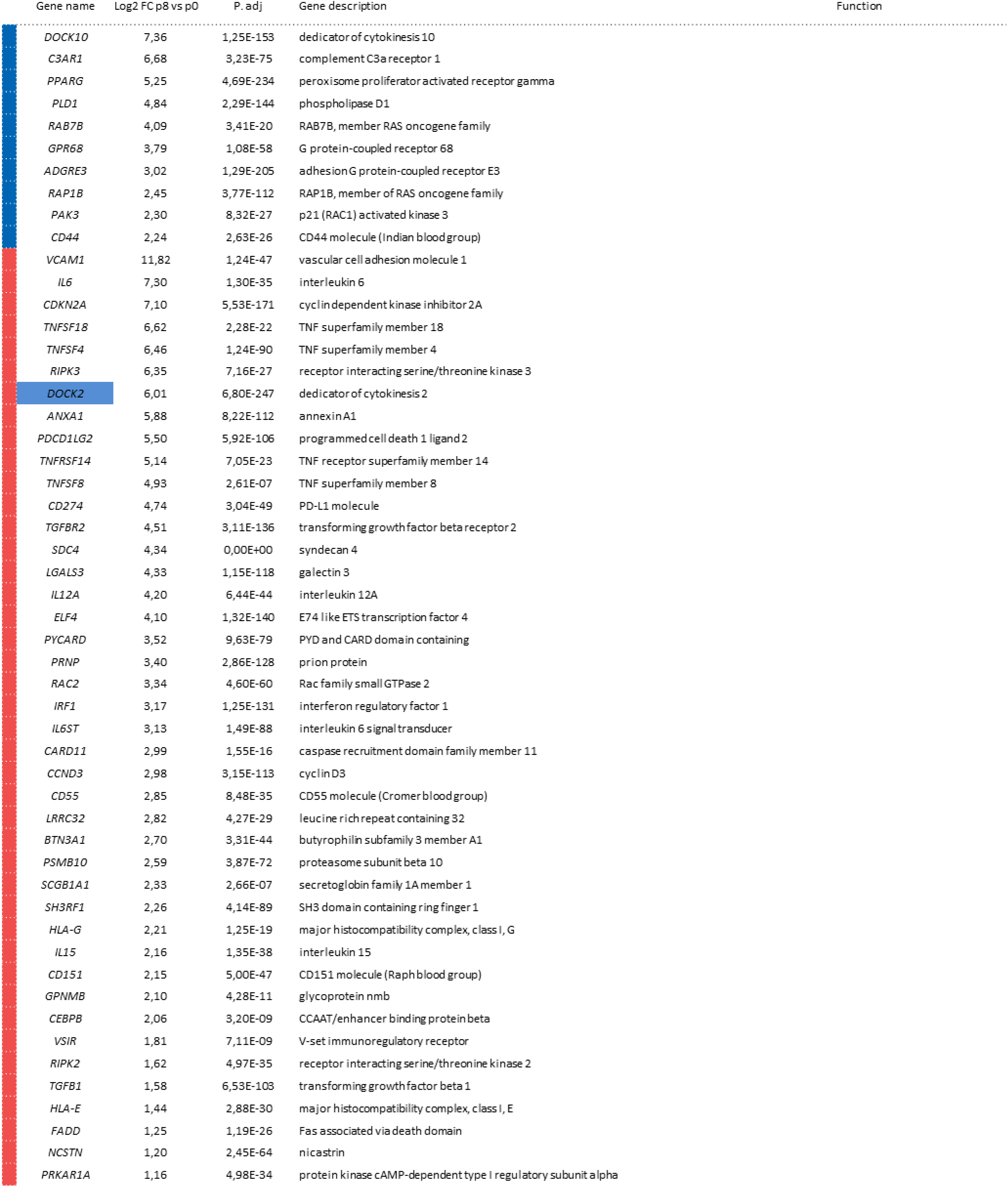
**Genes linked with T-cell proliferation (red) or G-protein signaling (blue)** found significantly upregulated in p8 iPS-MSCS compared to p0 iPS-MSCS and corresponding to parental MSCS. P values were adjusted using Benjamini-Hochberg correction.

**Figure S4a:**
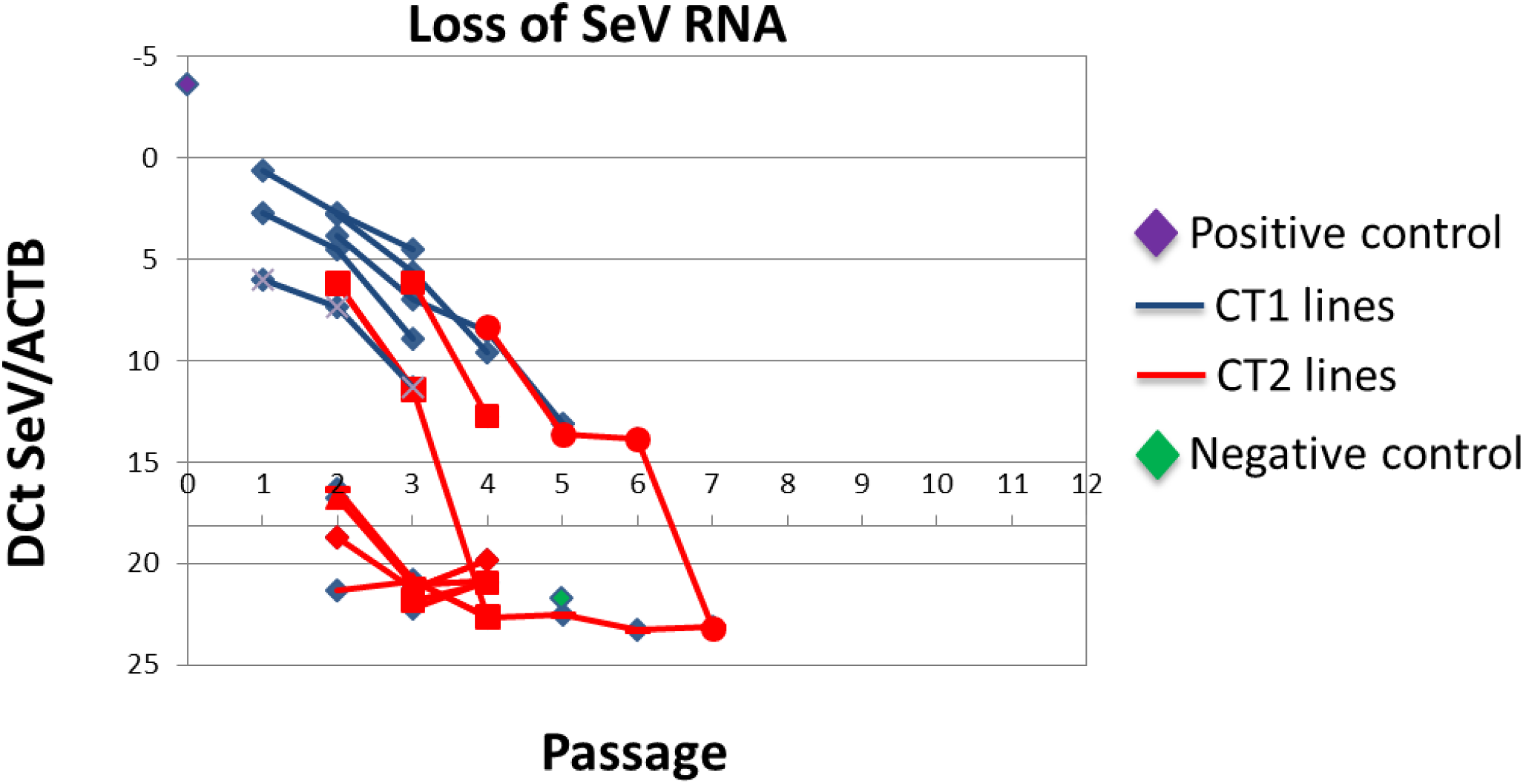
Characterization of iSPC lines. Single clones …

**Figure S4c:**
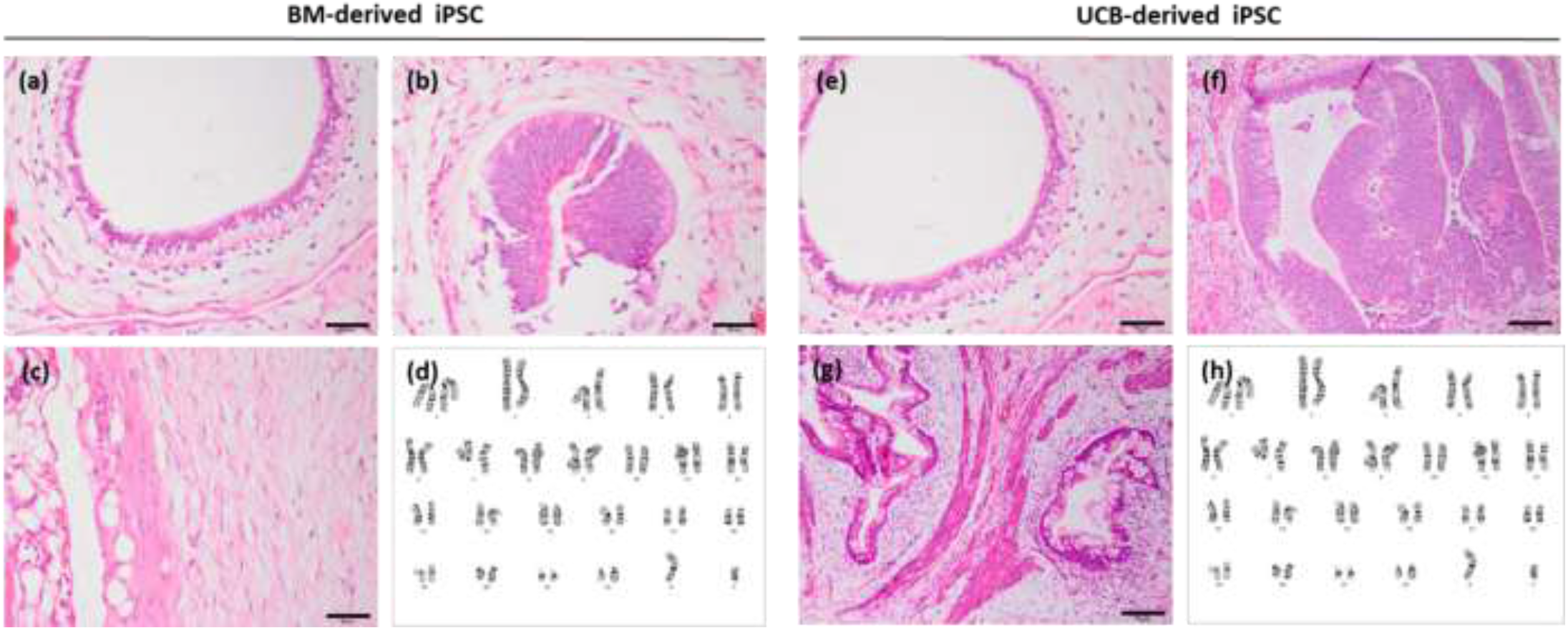
Characterisation of iPSC lines. Hematoxylin and eosin staining showing (**a**, **e**) endoderm (**b**, **f**) ectoderm (**c**, **g**) mesoderm structures in histology after teratoma test injection of iPSCs from BM- and UCB-derived MSCs as indicated. (**d**, **h**) Normal karyotype. Scale bars: (a, g) = 100 μm; (b, c, e, f) = 50 μm.

**Figure S4b:**
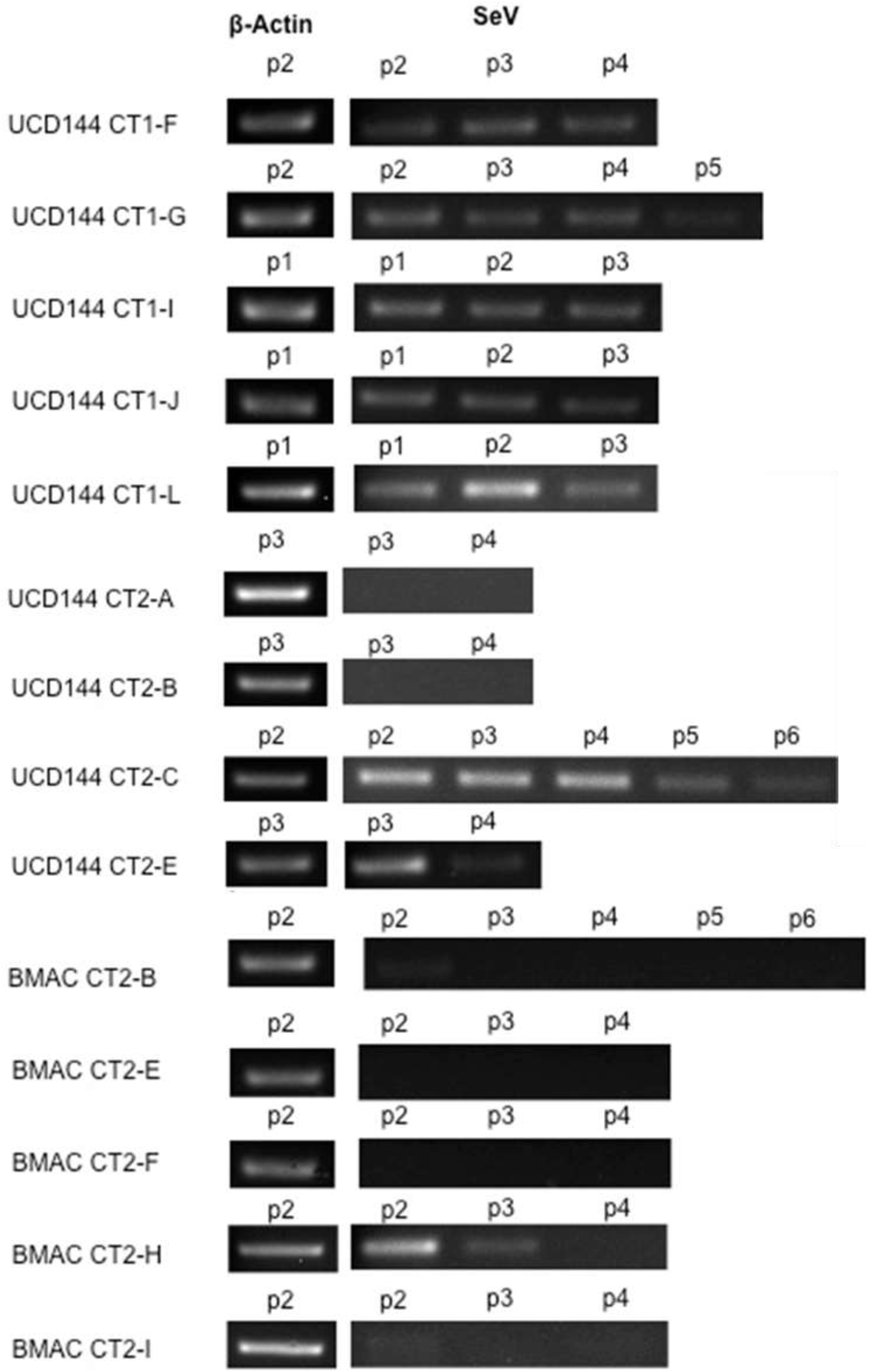
Characterization of iSPC lines. RTQ-PCR of single clones …cave: UCD reads UCB !!

**Figure S5:**
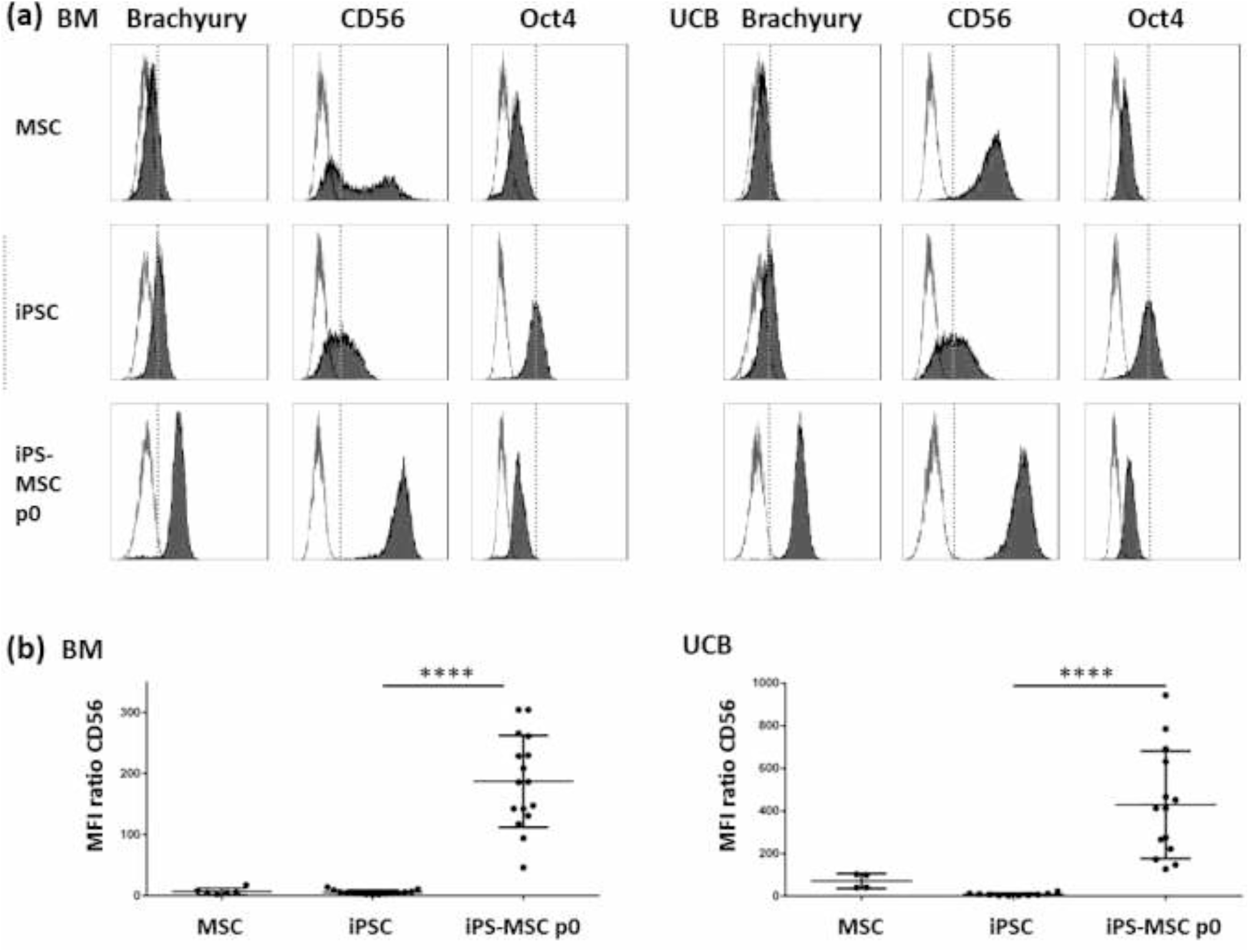
Mesoderm induction of iPS-MSCs at p0 was represented by the expression of Brachyury and CD56. (a) Representative histograms of flow cytometric analysis of MSCs, their iPSC progeny and mesoderm induced iPS-MSCs at p0 are depicted. Histograms show fluorescent cell surface (anti-CD56) and intracellular (anti-Brachyury and anti-Oct4) staining intensity of monoclonal antibodies conjugated to fluorophores (gray shading) and their corresponding isotype control (no shading). Increased brachyury and CD56 expression compared to iPS (dashed line) indicated mesoderm induction. Upon mesoderm induction, the expression of the pluripotency marker Oct4 decreased compared to iPS (dashed line). Histograms show the populations obtained following the hierarchical gating strategy: size and granularity, doublet exclusion, live cell population and corresponding marker. (b) Quantification of CD56 flow cytometric analysis revealed reproducible and significant induction of mesoderm in iPS-MSCs at p0 (**** p < 0.0001, n = 4-16).

**Figure S6:**
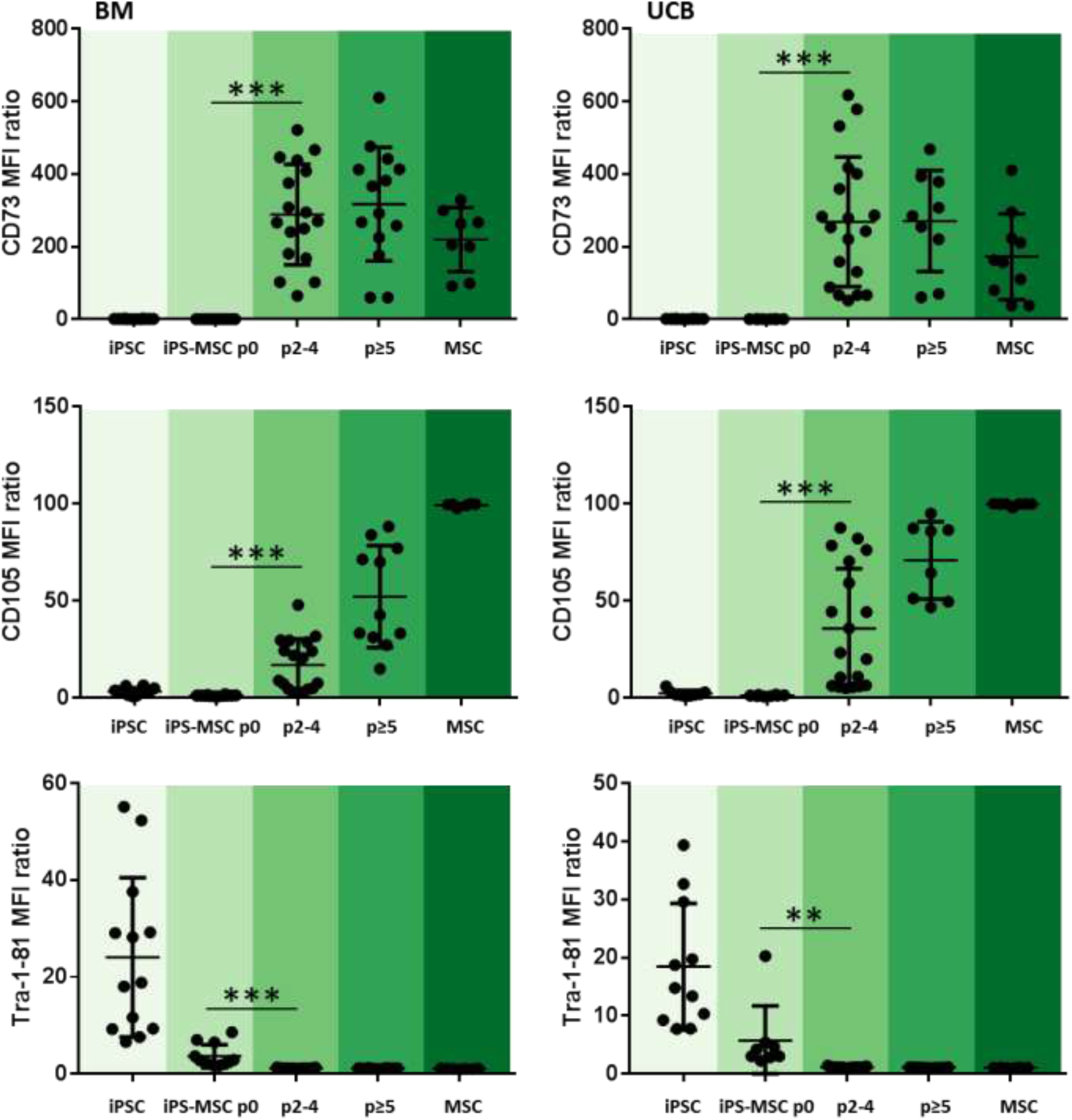
Monitoring of the differentiation of the MSC markers CD73 and CD105 and the pluripotency marker Tra-1-81 by flow cytometric analysis. The MSC markers increased from passage 2 onwards whereas Tra-1-81 expression was decreased already at passage 0 after mesoderm induction. Shown are MFI ratios of flow cytometric measurements (n = 6-19).

**Figure S7:**
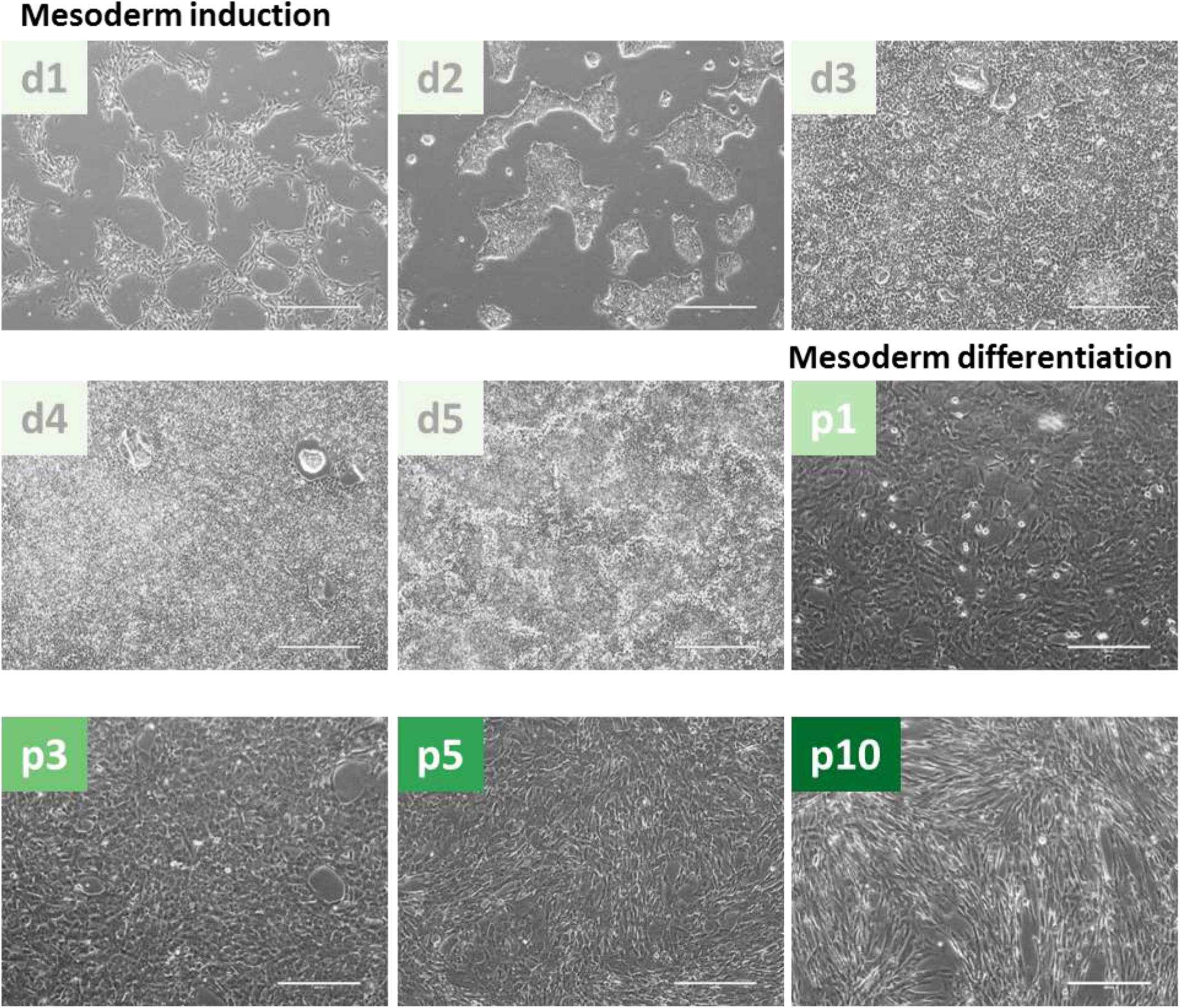
Mesoderm induction and differentiation of iPSCs with STEMdiff™ mesoderm induction medium and EGM-2/10% pHPL medium. Cells were monitored every day by microscopy after changing the medium. Shown are representative pictures of iPSCs derived from BM-MSCs (scale bar 400 μm).

**Figure S8:**
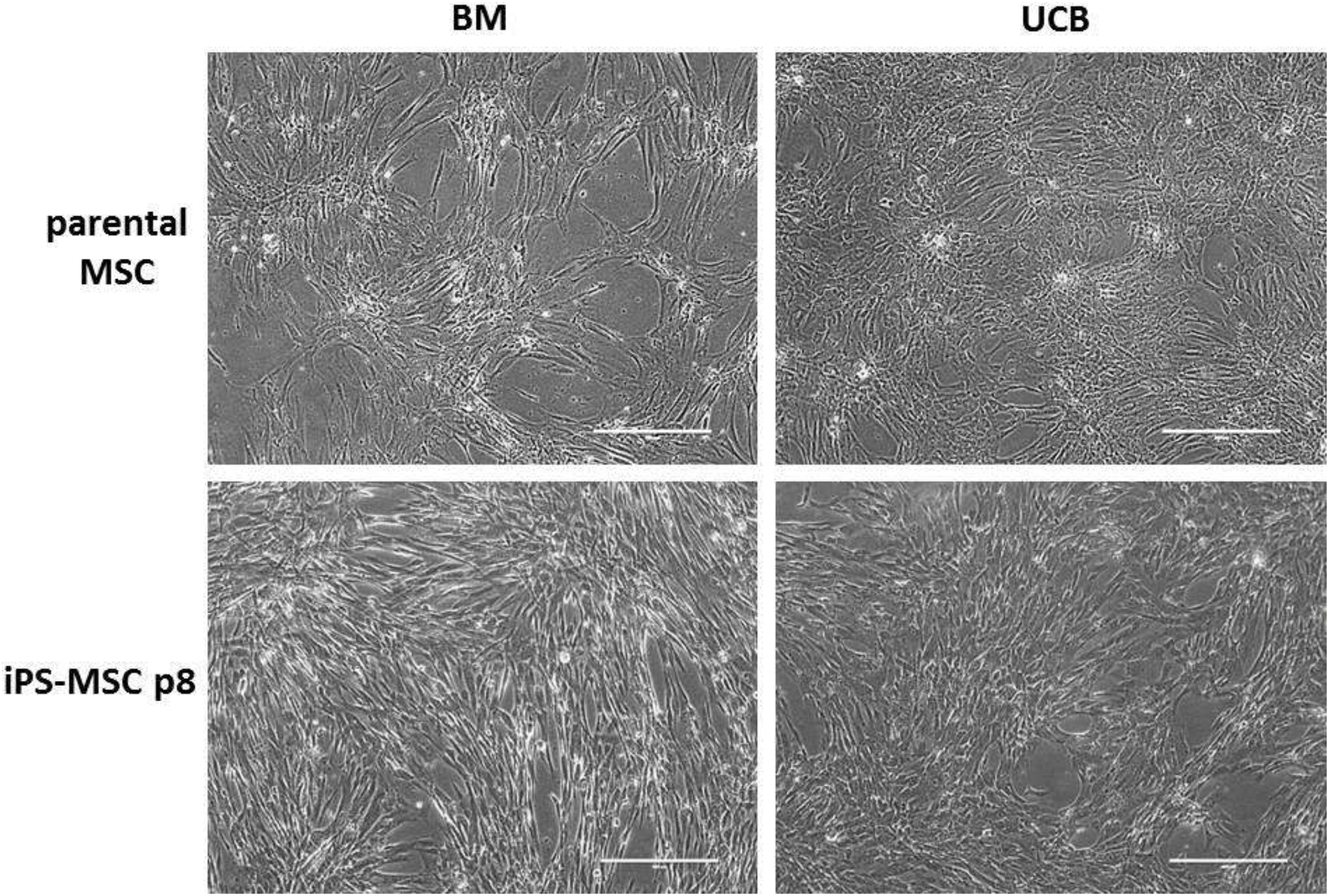
Morphology of iPS-MSCs. The morphology of iPS-MSCs was comparable to their parental MSCs. Shown are representative pictures of BM and UCB-derived MSCs and their iPS-MSC progeny (scale bar 400 μm).

**Figure S9:**
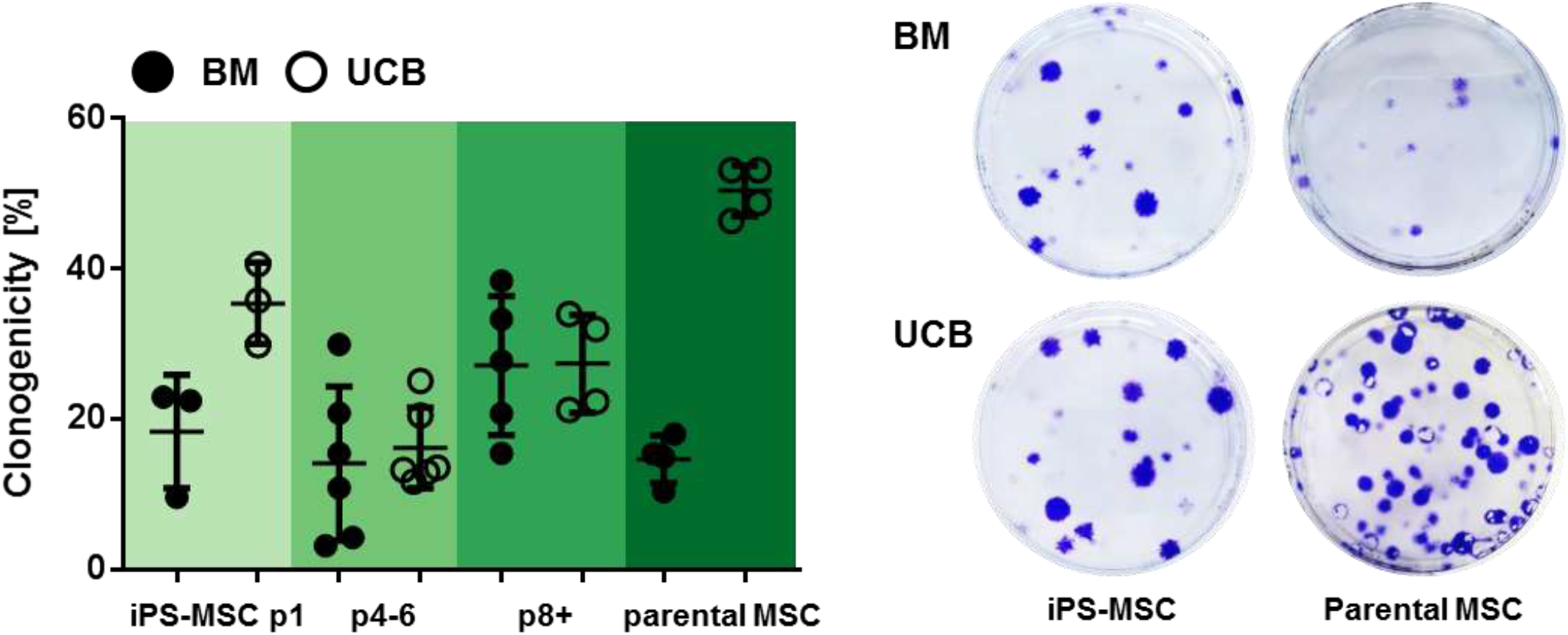
Clonogenicity of iPS-MSCs. Colony forming capacity (CFU-F) of iPS-MSCs compared to parental MSCs was assessed at a density of 3 cells per cm^2^ with ROCK inhibitor corresponding to the differentiation protocol from passage 1 to 5. Primary MSCs and iPS-MSCs passage ≥ 8 were seeded without ROCK inhibitor (n = 3-6).

**Figure S10:**
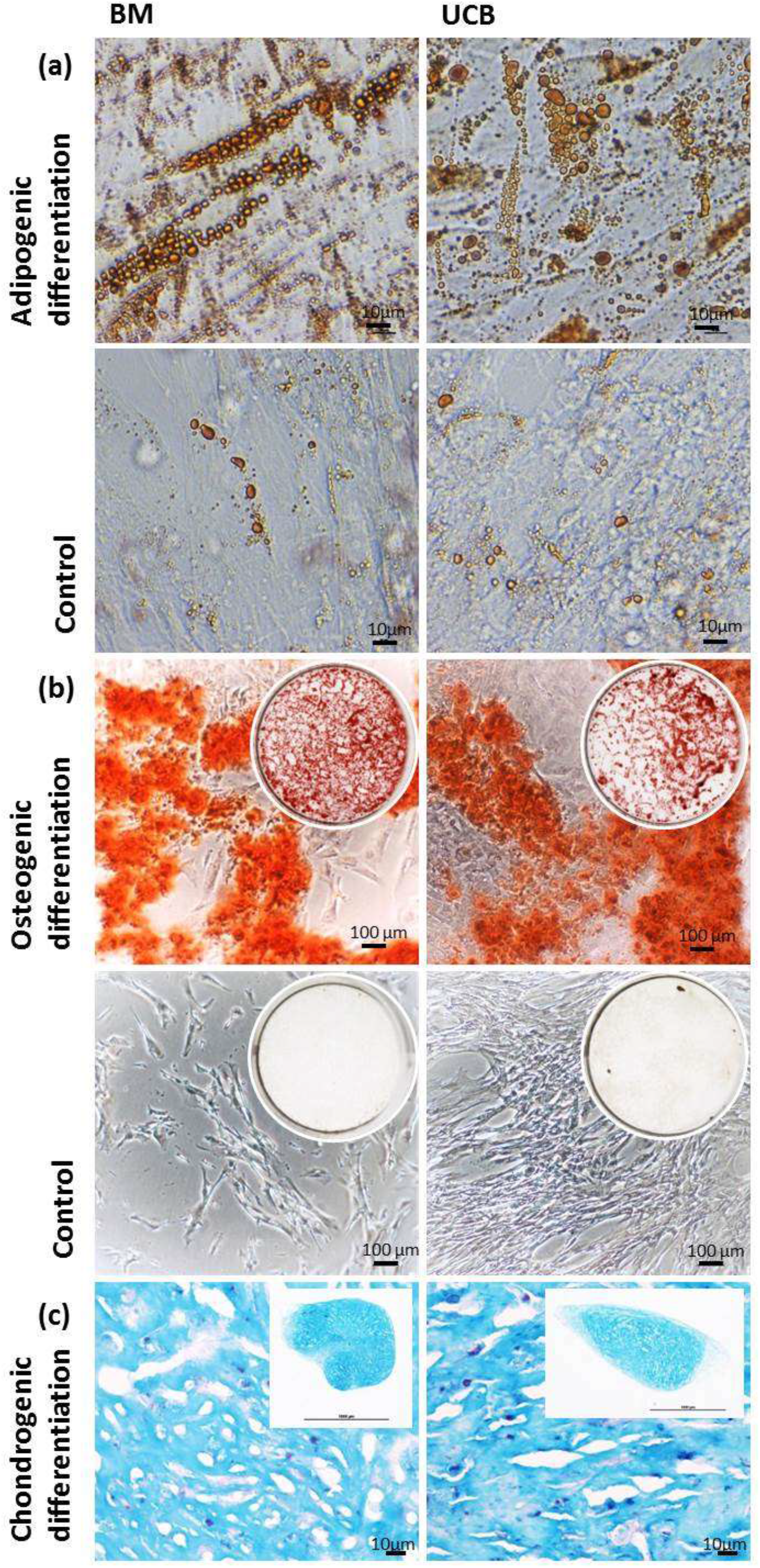
Adipogenic, osteogenic and chondrogenic differentiation potential of iPS-MSCs. Adipogenic and osteogenic differentiation potential of iPS-MSCs differentiated from bone marrow (BM) or umbilical cord blood (UCB)-derived iPS was revealed by positive Oil Red O (a) and Alizarin S Red (b) stainings respectively. Chondrogenic differentiation (c) was assessed by Alcian blue staining although the cells did not show the characteristic morphology of chondrocytes.

**Figure S11a:**
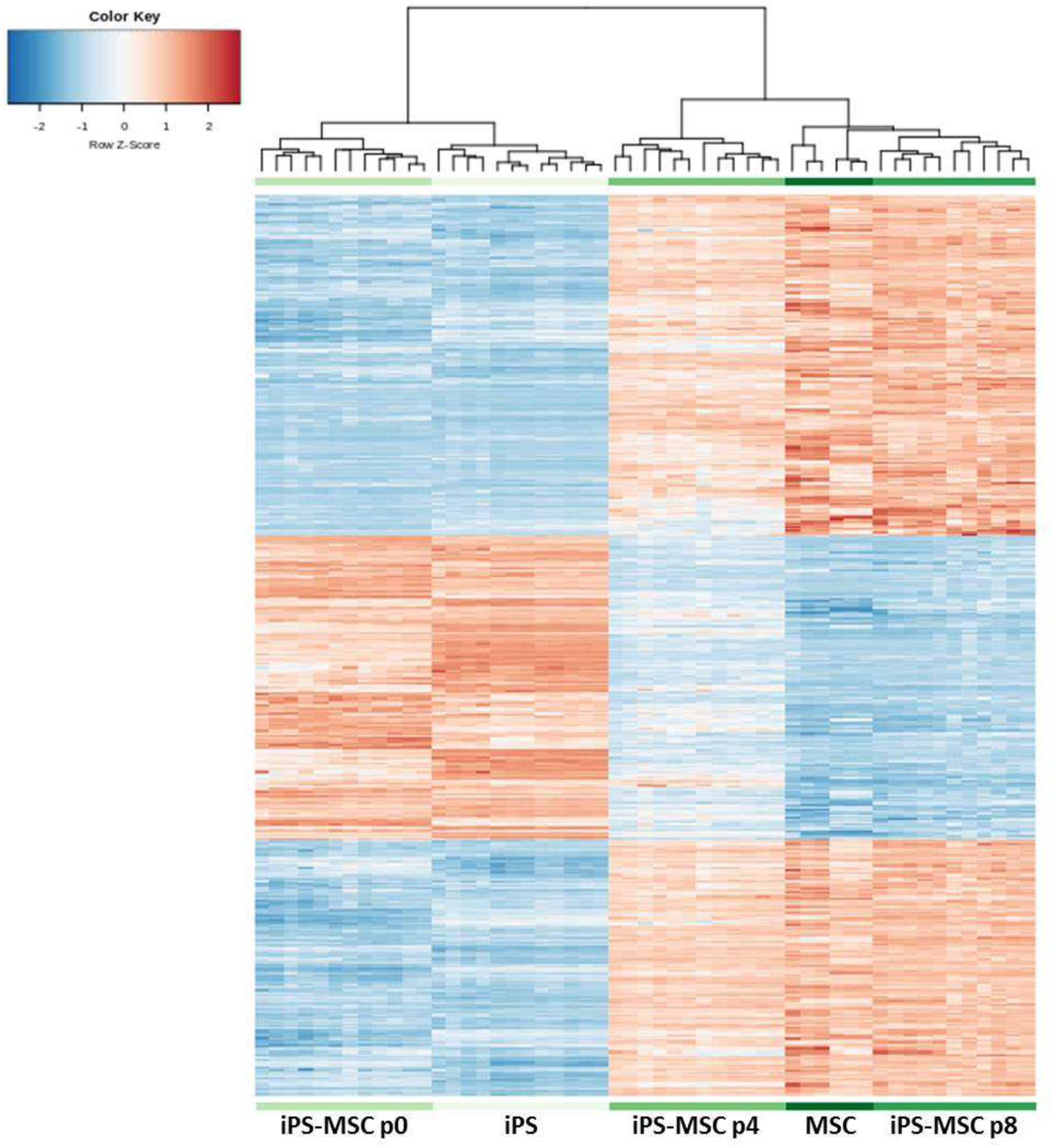
Hierarchical clustering heat map with expression values of the 500 most variable genes. Genes are shown in rows while the different samples are column listed. Side color bar on the top indicates the different cells or passages. Gene expression values were row Z-score normalized where lower expression is denoted by blue and higher expression by red color as shown in the legend above iPSCs (n = 12), p0 iPS-MSCs (n = 12), p4 iPS-MSCs (n = 12), p8 iPS-MSCs (n = 11), MSCs (n = 6).

**Figure S11b:**
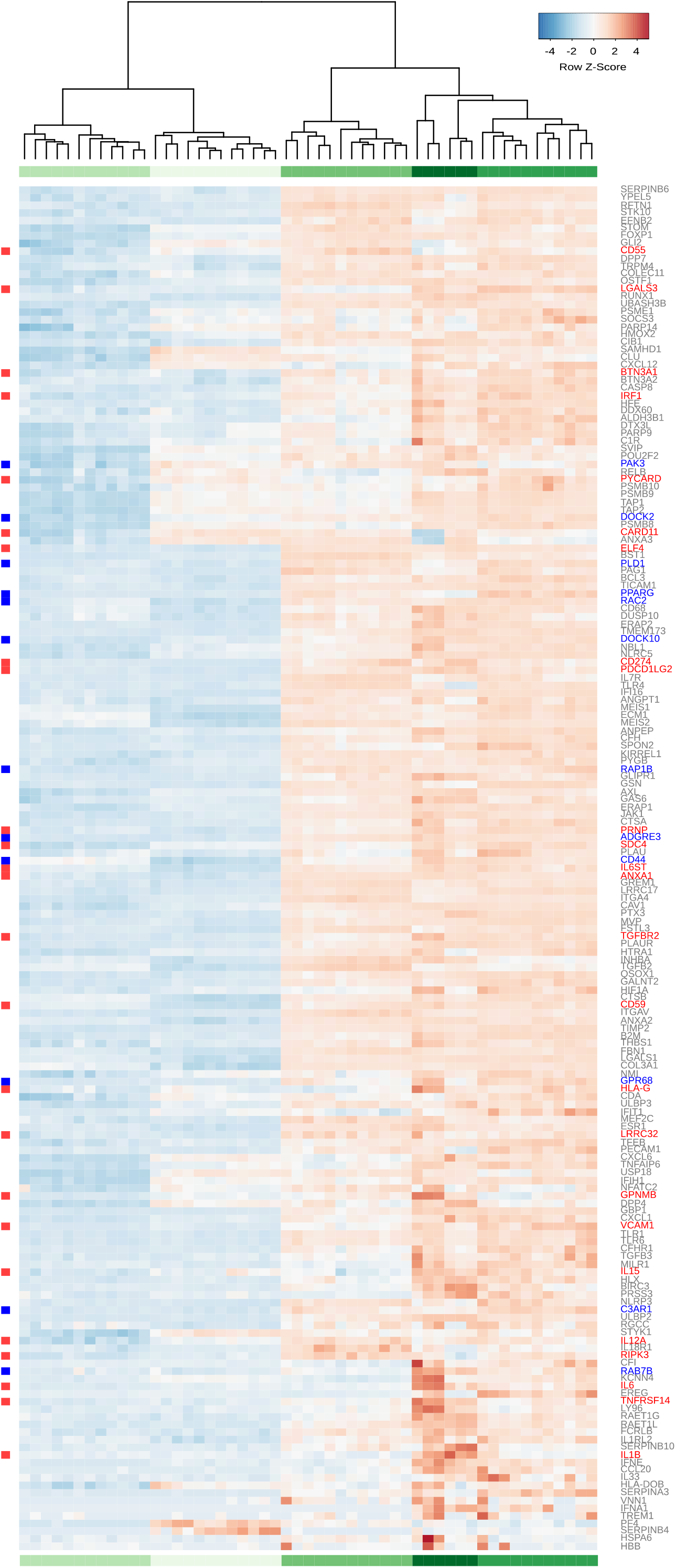
Alternative to S11a ….

**Figure S12:**
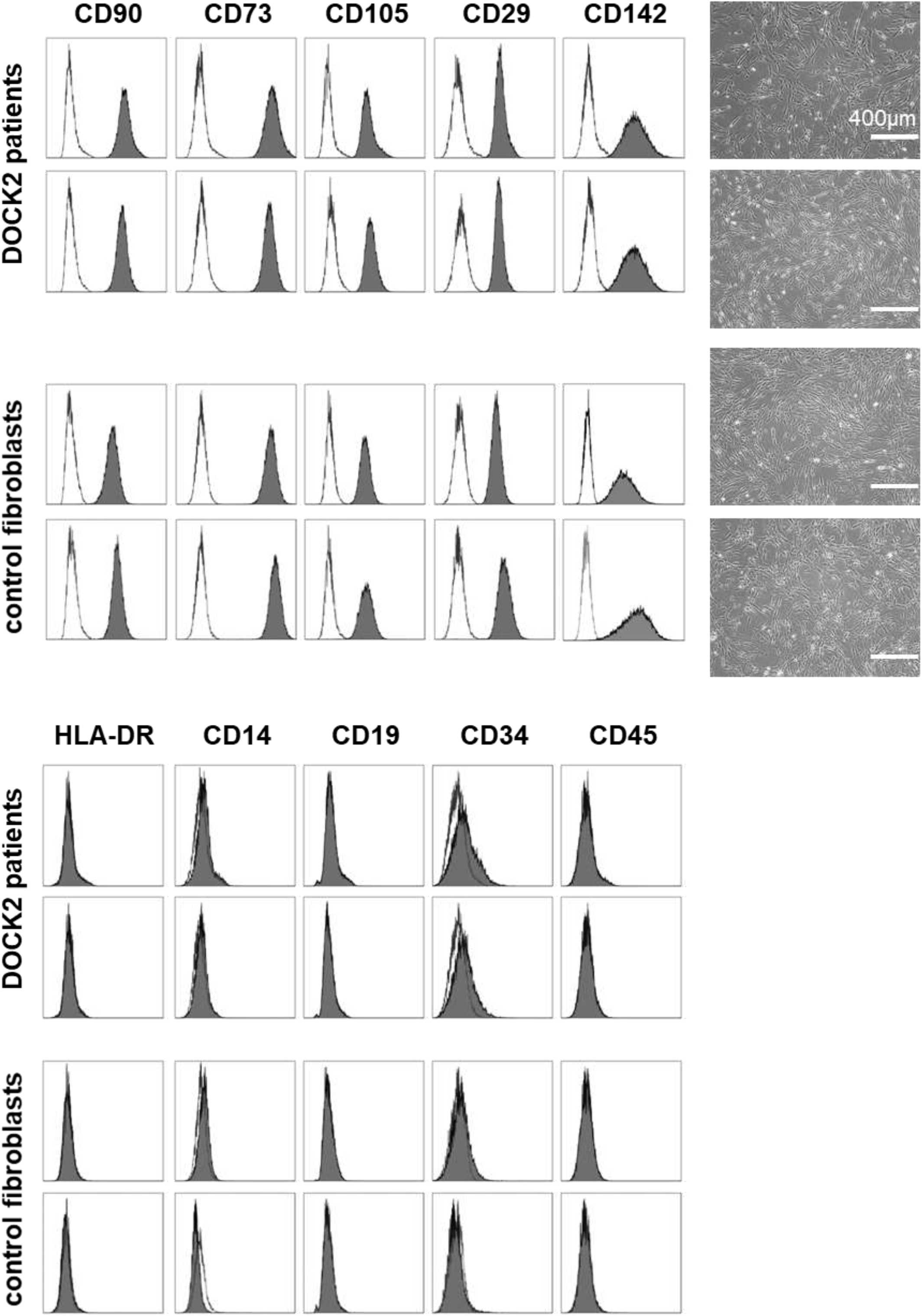
Immune phenotyping of fibroblasts derived from DOCK2 patients. Histograms show fluorescent cell surface staining intensity of anti-CD90, anti-CD73, anti-CD105, anti-CD142, anti-CD29, anti-HLA-DR, anti-CD14, anti-CD19, anti-CD34 and anti-CD45 monoclonal antibodies conjugated to fluorophores (gray shading) and corresponding isotype control (no shading). Morphology of DOCK2 patient and control fibroblasts by phase contrast microscopy, scale bar = 400 μm

**Figure S13:**
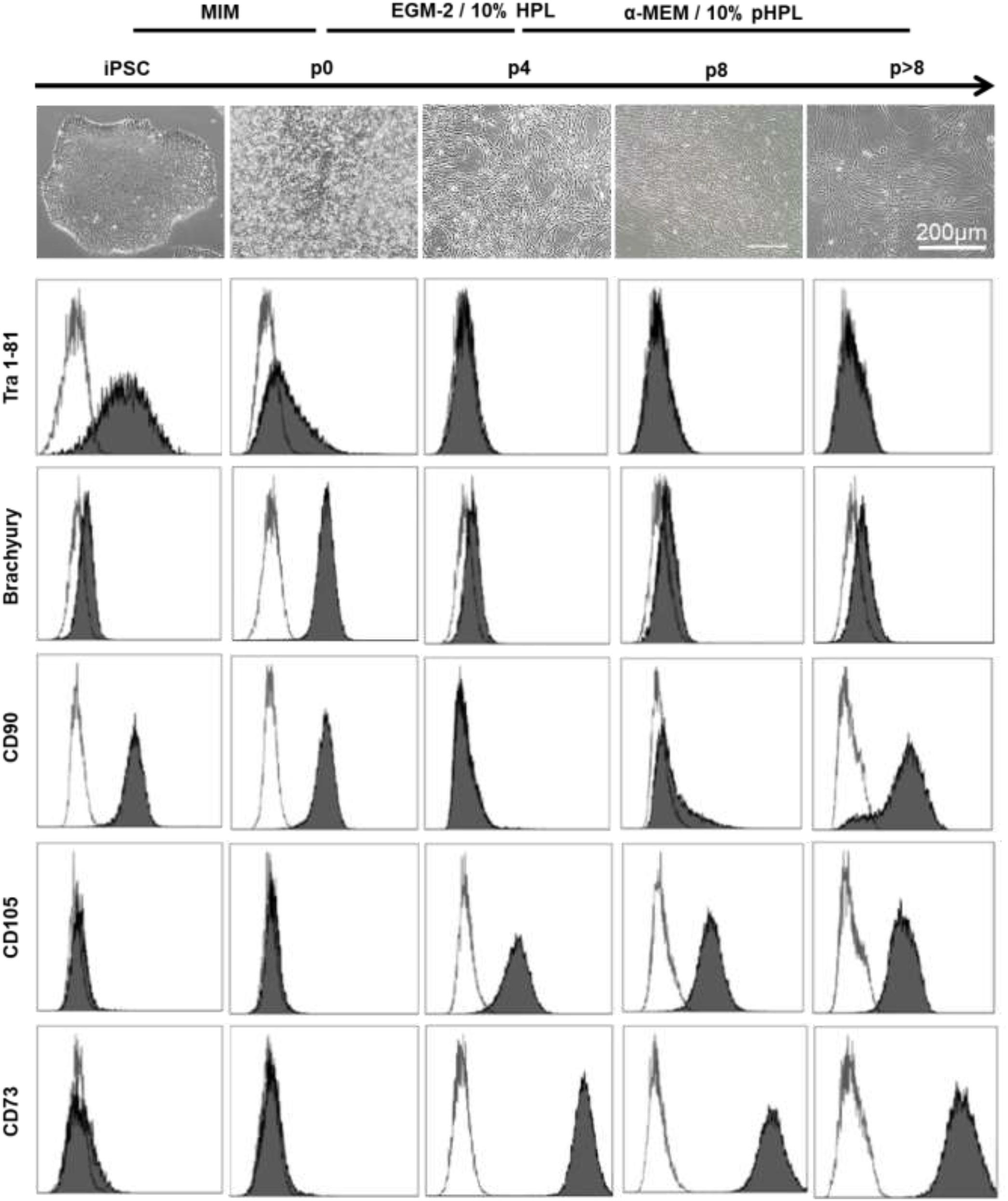
Final maturation of iPS-MSCs. An idditional maturation beyond passage 8 in another MSC-optimized medium …

**Figure S14:**
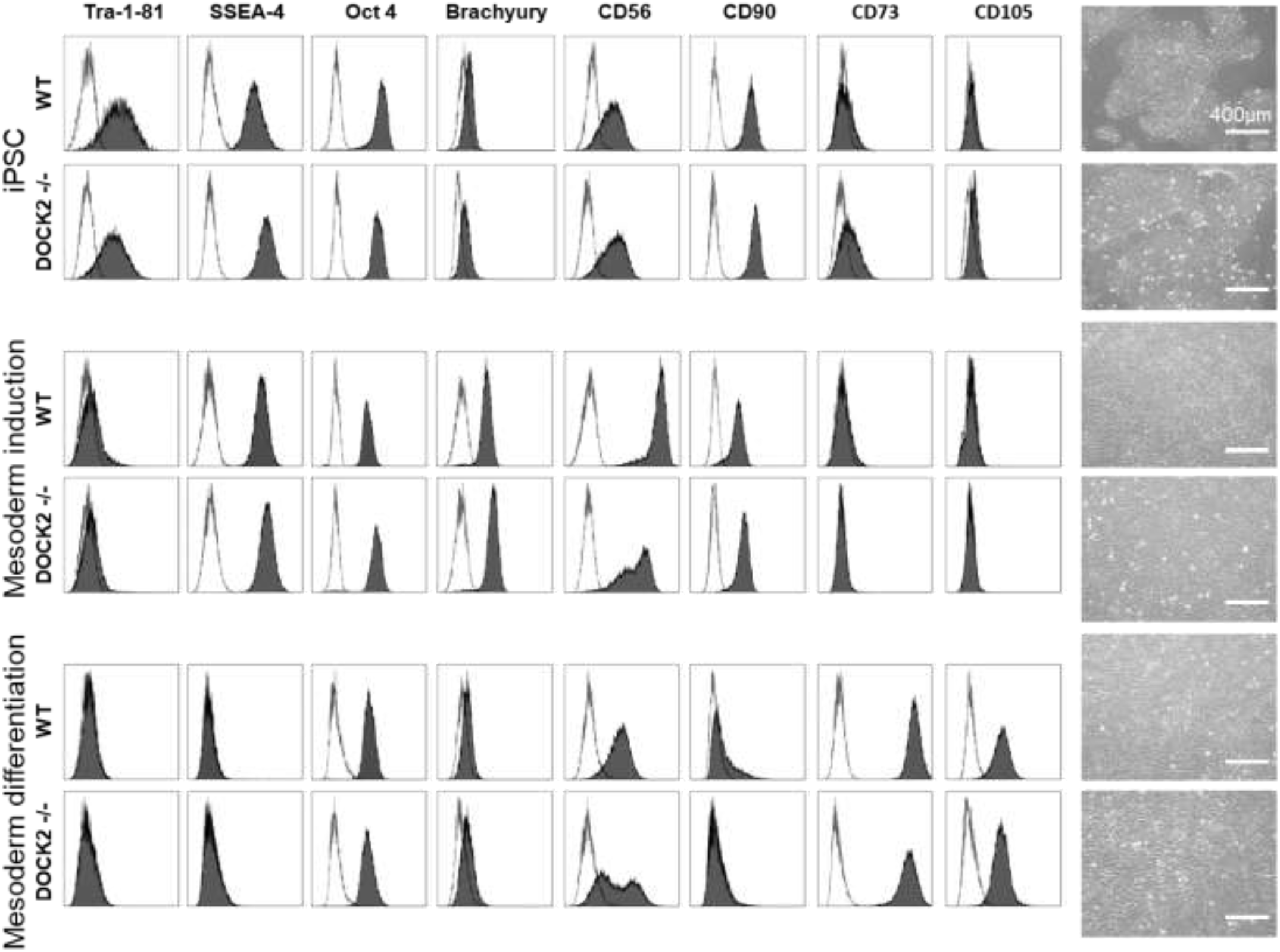
Marker expression profile after mesoderm induction and differentiation of DOCK2 knockout iPSC (CRISPR/Cas iPS-MSC DOCK2 −/−). Representative histograms of flow cytometric analysis of wildtype iPSC (wt, reprogrammed from UCB-derived MSC), wt iPSC-MSC after mesoderm induction and DOCK2 knockout (−/−) iPS-MSC (CRISPR/Cas-derived, one out of three clones is shown) and (b) their further differentiated iPS-MSC progeny are depicted. Histograms show fluorescent cell surface (anti-Tra 1-81, SSEA-4, CD56, CD90, CD73 and CD105) and intracellular (anti-Brachyury and anti-Oct4) staining intensity of monoclonal antibodies conjugated to fluorophores (gray shading) and their corresponding isotype control (no shading). Histograms show the populations obtained following the hierarchical gating strategy: size and granularity, doublet exclusion, live cell population and corresponding marker. Increased brachyury and CD56 expression compared to iPSC indicated mesoderm induction, although DOCK2 −/− iPS-MSC did not show a uniform CD56 expressing population (a). Mesoderm differentiated DOCK2 −/− iPS-MSC lost Tra-1-81and SSEA-4 expression and induced CD73 and CD105 comparable to wt iPS-MSC. The morphology of WT and DOCK2 −/− iPS-MSC was comparable (scale bar = 400 μm).

**Figure S15:**
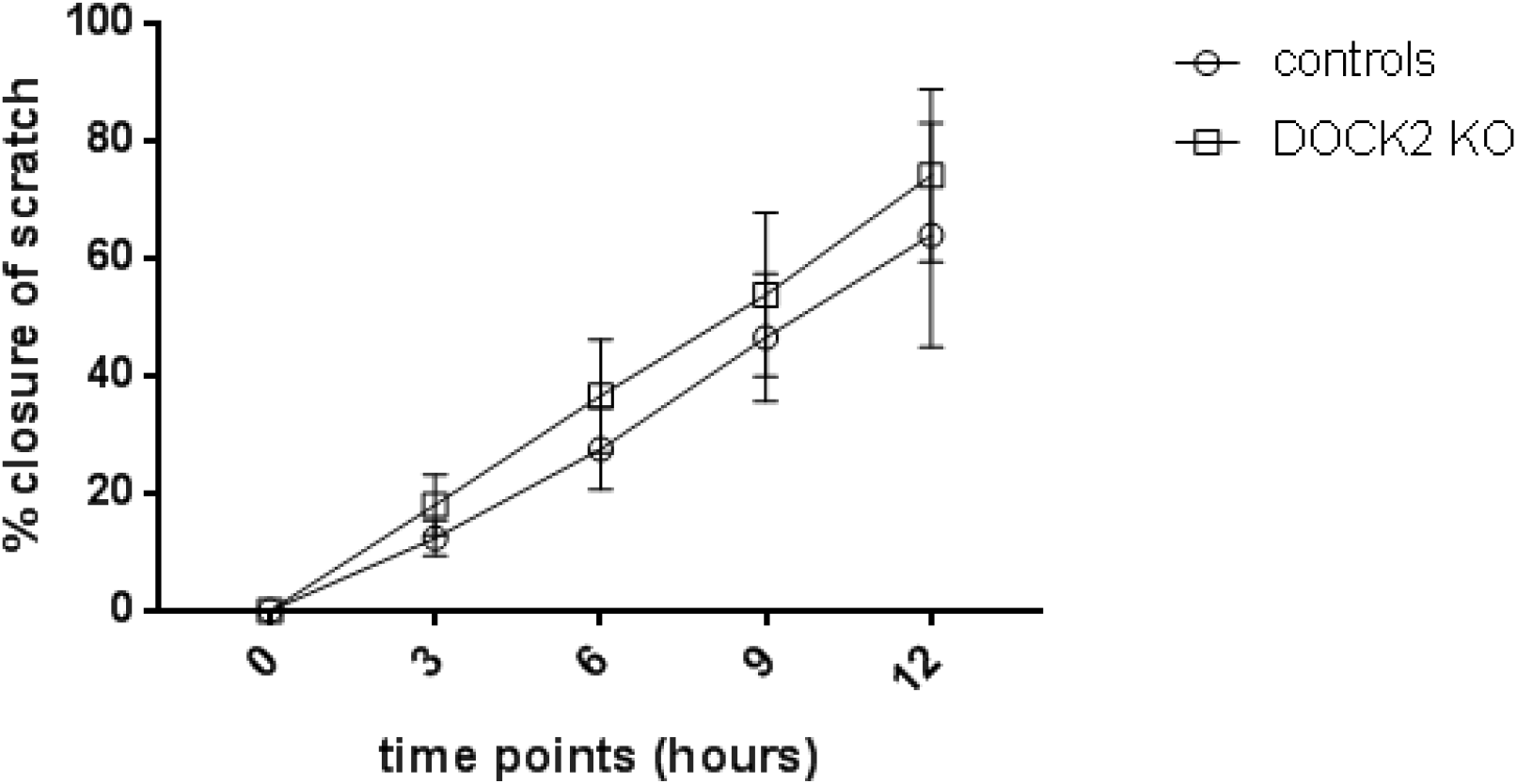
Scratch assay of fibroblasts derived from DOCK2 deficient patients vs. controls. Wounding a fibroblast monolayer in a scratch assay was used to test migratory wound repair of DOCK2 deficient fibroblasts compared to control fibroblasts. The assay was repeated three times with two different patients and two different control cells. DOCK2 deficient patient cells did not show an altered wound closure capacity.

